# Calcium determines *Lactiplantibacillus plantarum* intraspecies competitive fitness

**DOI:** 10.1101/2022.04.25.489480

**Authors:** Annabelle O. Yu, Lei Wei, Maria L. Marco

## Abstract

The importance of individual nutrients for microbial strain robustness and coexistence in habitats containing different members of the same species is not well understood. To address this for *Lactiplantibacillus plantarum* in food fermentations, we performed comparative genomics and examined the nutritive requirements and competitive fitness for *L. plantarum* strains B1.1 and B1.3 isolated from a single sample of teff injera fermentation batter. Compared to B1.1 and other *L. plantarum* strains, B1.3 has a smaller genome, limited biosynthetic capacities, and large mobilome. Despite these differences, B1.3 was equally competitive with B1.1 in a suspension of teff flour. In commercially-sourced, nutrient-replete MRS (cMRS) medium, strain B1.3 reached three-fold higher numbers than B1.1 within two days of passage. Because B1.3 growth and competitive fitness was poor in mMRS, a modified MRS lacking beef extract, we used mMRS to identify nutrients needed for robust B1.3 growth. No improvement was observed when mMRS was supplemented with nucleotides, amino acids, vitamins, or monovalent metals. Remarkably, the addition of divalent metal salts increased the growth rate and cell yields of B1.3 in mMRS. Metal requirements were confirmed by Inductively Coupled Plasma Mass Spectrometry, showing that total B1.3 intracellular metal concentrations were significantly (up to 2.7-fold) reduced compared to B1.1. Supplemental CaCl_2_ conferred the greatest effect, resulting in equal growth between B1.1 and B1.3 over successive five passages in mMRS. Moreover, calcium supplementation reversed a B1.3 strain-specific stationary phase, flocculation phenotype. These findings show how *L. plantarum* calcium requirements affect competitive fitness at the strain level.

**Importance:** Ecological theory states that the struggle for existence is stronger between closely related species. Contrary to this assertion, fermented foods frequently sustain conspecific individuals, despite their high levels of phylogenetic relatedness. Therefore, we investigated two isolates of *Lactiplantibacillus plantarum* B1.1 and B1.3 randomly selected from a single batch of teff injera batter. These strains spanned the known genomic and phenotypic range of the *L. plantarum* species, and in nutrient-replete, laboratory culture medium, strain B1.3 exhibited poor growth and was outcompeted by the more robust strain B1.1. Despite those differences, B1.1 and B1.3 were equally competitive in teff flour. This result shows how these bacteria have adapted for co-existence in that environment. The capacity for the single macronutrient calcium to restore B1.3 competitive fitness in laboratory culture medium suggests that *L. plantarum* intraspecies diversity found in food systems is fine-tuned to nutrient requirements at the strain level.

## Introduction

Lactic acid bacteria (LAB) are essential for the production of thousands of different types of fermented dairy, meat, and plant-based foods and beverages (1), where they can account for over 95% of all microbes present (2, 3). These bacteria are also inhabitants of the human, animal, and insect microbiomes (4). A common feature among LAB-associated habitats is that they are nutrient-rich, frequently containing high quantities of sugars as well as other macro- and micro- nutrients (e.g., vitamins, amino acids, trace metals, and nucleotides). Consistent with this assertion, numerous studies have shown that LAB strains have undergone genomic reduction as a result of adapting to the nutrient-rich environments in which they are found (e.g. food fermentations and mucosal surfaces) (5–9). Gene gain and loss likely occurs as a function of the selective pressures occurring in food habitats (e.g. yogurt (10)). However, the extent of intraspecies variation among LAB within food fermentations and the consequences of strain differences on ecological robustness remain to be determined.

*Lactiplantibacillus plantarum* (formerly known as *Lactobacillus plantarum*) is one of the most frequently isolated LAB species from fresh and fermented foods (vegetables, fruits, dairy, and meat) and intestinal sources (11–13). Because of its capacity to colonize different plant and animal habitats and its genome plasticity, *L. plantarum* is regarded to be a “generalist” or “nomadic” LAB (14–16). Strains of *L. plantarum* tend to have larger genome sizes (3 to 3.6 Mbp) compared to other LAB (1.5 to 3.0 Mbp) and fewer nutritional requirements (14, 17). However, as shown for the model strain WCFS1, *L. plantarum* still requires numerous vitamins (D-pantothenate, D-biotin, nicotinic acid, and riboflavin) and amino acids (arginine, glutamic acid, glycine, leucine, methionine, tryptophan, and valine) for robust growth (18).

Recent comparative genomic analysis and phenotypic screening approaches have shown the tremendous intraspecific variation in the *L. plantarum* species (14, 19, 20). We recently reported that *L. plantarum* from the same (fermented) plant food type tend to use the same carbohydrates for growth and exhibit similar stress tolerance levels, indicating the presence of strong selective pressures for strain selection and adaptation within those environments (20). However, despite certain similarities, strain variation is not limited to isolation source, and we and others showed that *L. plantarum* strains recovered from the same food, plant, or animal environment can have very different genotypic and phenotypic properties (19). Although it is feasible that this conspecific diversity is important for sustaining *L. plantarum* within individual habitats, the specific traits needed for sustaining strain co-existence should be identified.

*L. plantarum* B1.1 and B1.3 were isolated from a single sample of teff injera and were previously compared to each other and to other plant-associated *L. plantarum* strains (Yu *et al*., 2021). The carbohydrate utilization and stress tolerance capacities of strain B1.1 were robust and similar to reference strain NCIMB8826R and other *L. plantarum* isolates. Conversely, growth of strain B1.3 was poor on glucose and other sugars and was also unable to tolerate acidic pH, and high salt, ethanol, and high temperature conditions (20). To determine how strain B1.3 could be competitive in teff injera fermentations where other *L. plantarum* strains such as B1.1 are present, we investigated B1.3 and B1.1 for their genome characteristics, competitive fitness, and specific nutritional requirements for robust growth in nutrient-replete MRS laboratory culture medium. These studies led us to identify how the requirement for divalent metal cations and calcium specifically differs between strains of *L. plantarum* and how those nutrients are important for sustaining B1.1 and B1.3 in co-culture.

## Results

### *L. plantarum* B1.3 contains a large mobilome and lacks numerous biosynthetic pathways present in other *L. plantarum* strains

The genome of *L. plantarum* B1.3 (3.09 Mbp) is approximately 111 Kbp smaller than the genome of strain B1.1 (3.17 Mbp) (20). Compared to 663 *L. plantarum* genome assemblies, the genome size of strain B1.3 ranks in the lowest 10%. Consistent with its smaller size, the genome of *L. plantarum* B1.3 contains deletions ranging in size from 2,200 to 25,000 bp at different locations in the chromosome relative to the reference strain *L. plantarum* WCFS1 (**Table S1**). These deletions frequently occur in gene loci coding for amino acid and vitamin biosynthetic pathways and transporters and for the production of cell-surface associated and carbohydrate utilization proteins (**Table S1**). For example, B1.3 lacks genes required for the synthesis of thiamin, riboflavin, cystine, aromatic amino acids, and branched chain amino acids (**Table S1 and Fig. S1**). By comparison, the genome composition of strain B1.1 is similar to other plant-associated *L. plantarum* strains and the model *L. plantarum* strain WCFS1 (21, 22) .

The number of predicted proteins in individual COG categories also differed between *L. plantarum* B1.3 and B1.1 (**Table 1**). Strain B1.1 contains higher numbers of gene clusters relative to B1.3 in the transcription (K), energy production and conversion (C), nucleotide (F), coenzyme (H), lipid (I), and inorganic ion (P) transport and metabolism COGs (**Table 1**). Strain B1.3 has more gene clusters than B1.1 for cell wall/membrane/envelope biogenesis (M), post-translation modification/protein turnover/chaperone (O), replication/recombination/repair (L), and amino acid transport and metabolism (E) COGs. The greatest difference between the two strains however is in the mobilome COG. Strain B1.3 is annotated to contain 346 gene clusters, whereas B1.1 contains only 118, constituting nearly a three-fold difference (**Table 1**). Over 90% of B1.3 genes in the mobilome COG are annotated as transposases and the remainder are predicted to encode prophages or have a role in plasmid replication.

**Table 1.**
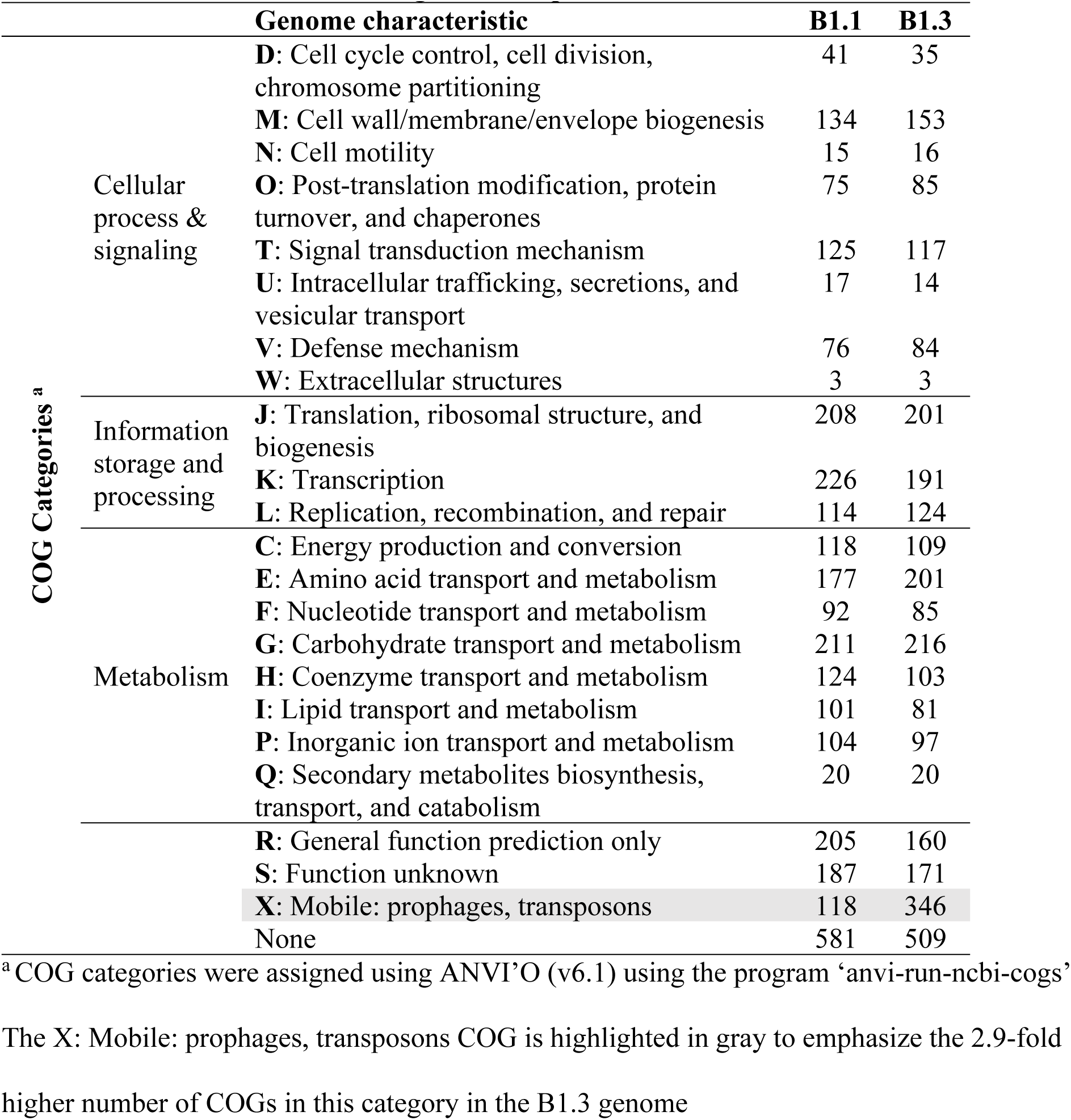
Distribution of COG categories in *L. plantarum* B1.1 and B1.3 Genome characteristic B1.1 B1.3.

### *L. plantarum* B1.3 growth is restored in mMRS containing beef extract

Consistent with the reduced biosynthetic capacity of B1.3, that strain grew poorly in mMRS, a modified version of MRS medium lacking beef extract (**Fig. 1**) (see also (20)). B1.3 exhibited biphasic growth in mMRS with lower growth rates (0.38 h^-1^ ± 0.01 and then 0.07 h^-1^ ± 0.01) compared to strain B1.1 (0.61 h^-1^ ± 0.01) (**Fig. 1**). Additionally, it was observed that within the first 24 h after stationary phase was reached, B1.3 but not B1.1 flocculated, forming aggregates (biofilm) which deposited on the bottom of the test tube (**Fig. 2**).

**Fig. 1.**
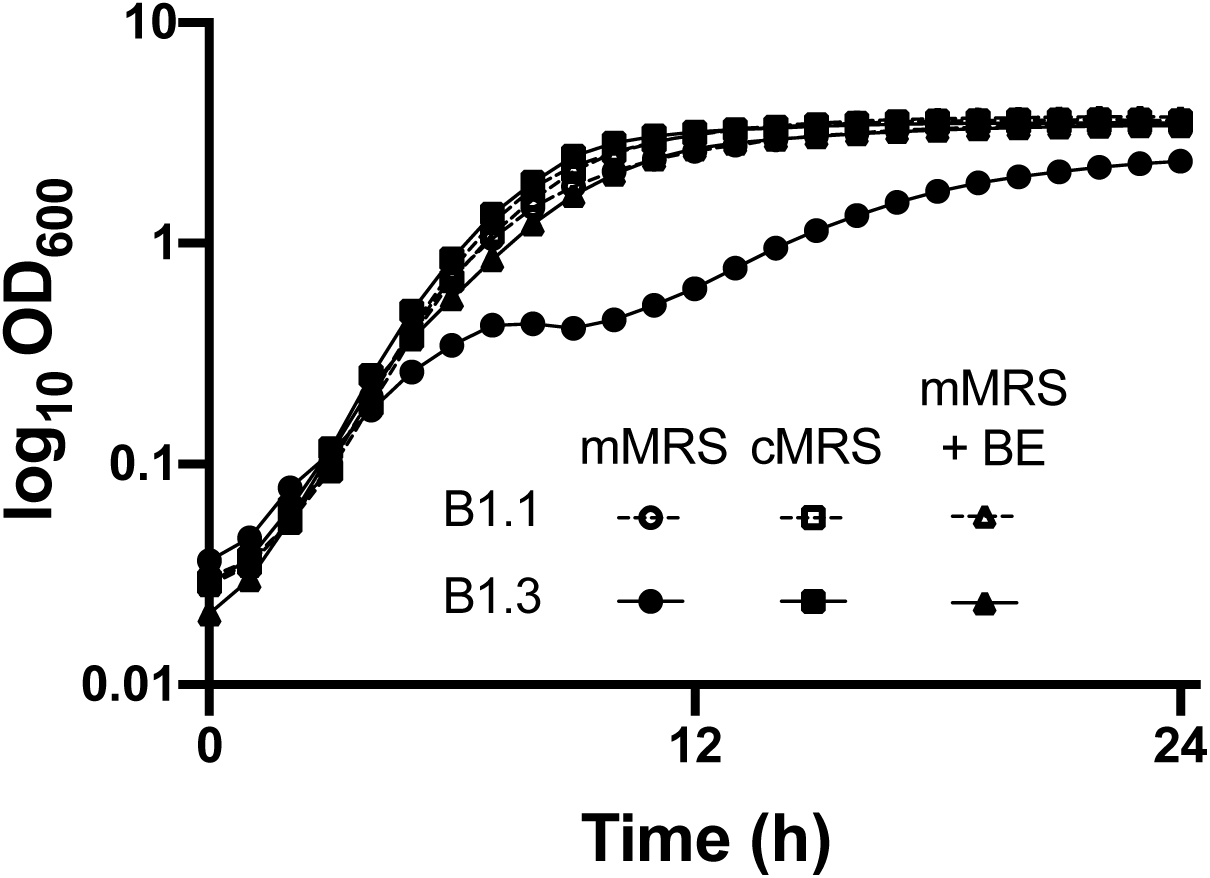
Growth of *L. plantarum* B1.3 is restored in mMRS supplemented with beef extract. *L. plantarum* B1.1 and B1.3 were inoculated into mMRS, cMRS, or mMRS supplemented with beef extract ((BE) 8 g/L) and incubated at 30 °C for 24 h. The avg ± stdev OD_600_ of three replicate cultures is shown.

**Fig. 2.**
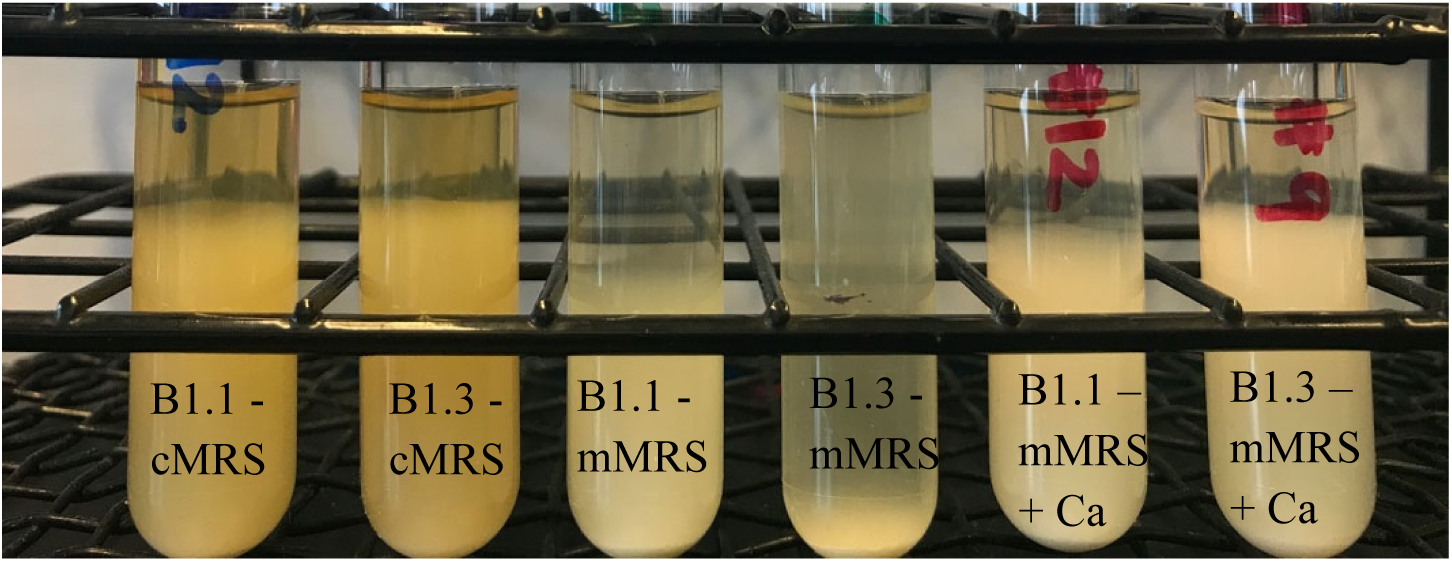
Auto-aggregation of *L. plantarum* B1.1 and B1.3 in cMRS, mMRS, and mMRS supplemented with 5mM CaCl_2_. *L. plantarum* B1.3 and B1.1 cultures were imaged after incubation at 30 °C for 72 h.

Notably, these differences between B1.1 and B1.3 were not found when the strains were incubated in a commercially sourced MRS (cMRS) containing beef extract (**Fig. 1 and Fig. 2**). The growth rate of B1.3 in cMRS (0.61 h^-1^ ± 0.01) was not significantly different from B1.1 (0.61 h^-1^ ± 0.01) in the cMRS laboratory culture medium, nor was there evidence of flocculation (**Fig 2**). The importance of beef extract (BE) for B1.3 growth in cMRS was confirmed by supplementing BE into mMRS (mMRS-BE) (**Fig. 1**). In mMRS-BE, the growth rate of B1.3 (0.62 h^-1^ ± 0.01) was equal to B1.1 (0.63 h^-1^ ± 0.01) and to both strains in cMRS. Furthermore, with the addition of BE, aggregation of B1.3 was no longer observed and instead matched the profile seen when B1.3 was grown in cMRS (**Fig. S2**). These findings show that the robustness of B1.3 growth is dependent on a factor provided by the BE.

### Strain B1.3 outcompetes B1.1 in cMRS and teff flour

We next determined the competitive fitness of B.1.3 and B1.1 in co-inoculation assays in which the strains were inoculated in equal proportions and passaged daily in fresh culture medium for five consecutive days. As expected, based on growth rates, strain B1.3 was rapidly outcompeted by B1.1 in mMRS (**Fig. 3A**). B1.3 was no longer detectable (detection limit of 10^7^ CFU/mL) within 24 h incubation in mMRS. Cell numbers of strain B1.1 increased from 1 x 10^5^ CFU/ml to 2 x 10^9^ CFU/ml during that time, reaching at least a 100-fold increase over B1.3 (**Fig. S3**). Remarkably, B1.3 outcompeted B1.1 in cMRS (**Fig. 3B**) and in mMRS-BE (**Fig. 3C**). Within two days of sequential passage in cMRS, strain B1.3 outgrew B1.1 in a ratio of approximately three to one (**Fig. 3B**). Strain B1.3 reached higher numbers than B1.1 in mMRS-BE as well, although the increases were not significant (p < 0.05, Student T’s-test) (**Fig. 3B and Fig. S3**).

**Fig. 3.**
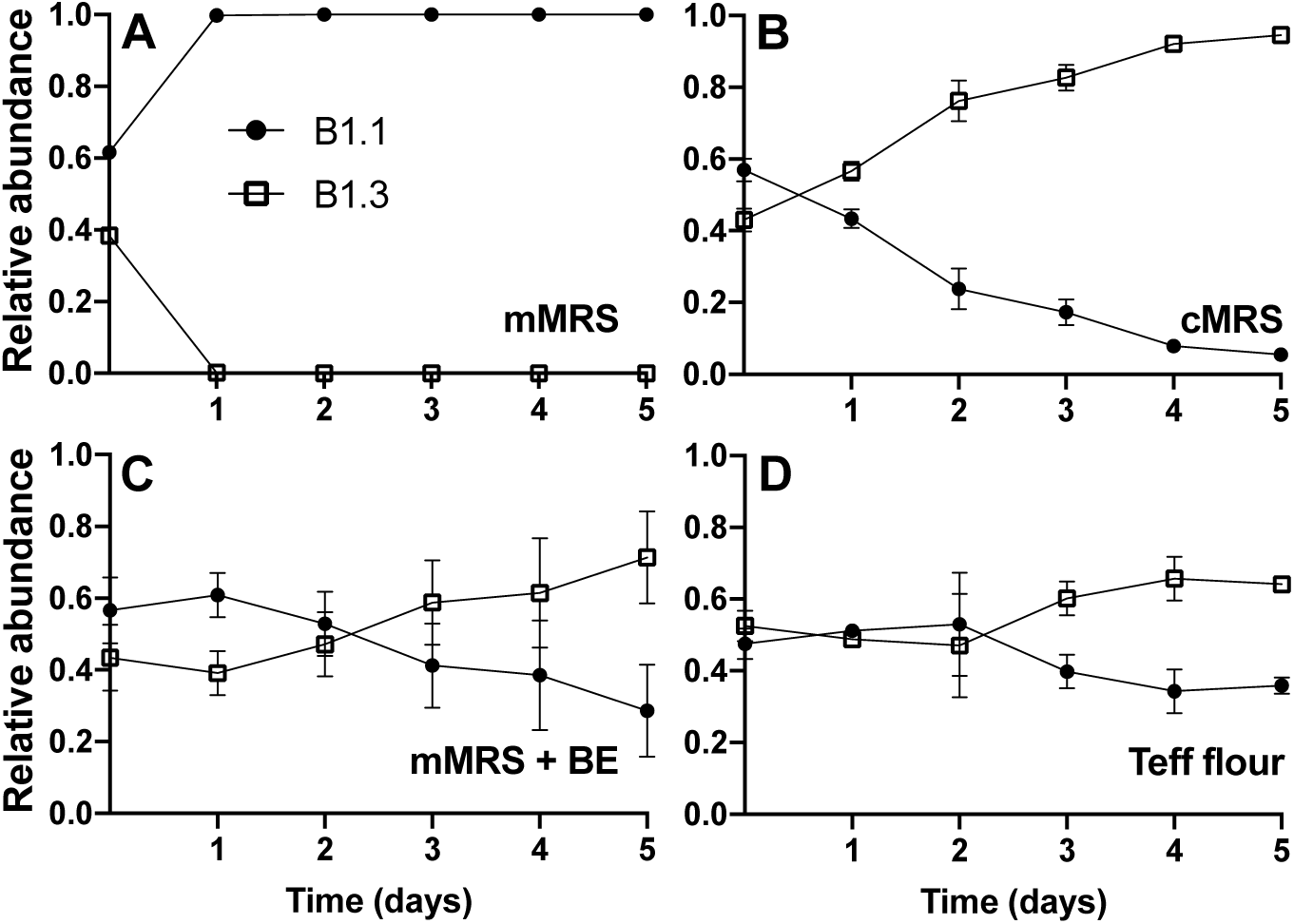
*L. plantarum* B1.3 exhibits a high level of competitive fitness compared to B1.1 in mMRS with beef extract (BE) and in teff flour. Equal numbers of *L. plantarum* B1.1 and B1.3 (10^5^ CFU/ml) were co-inoculated in mMRS, cMRS, mMRS supplemented with beef extract ((BE) 8 g/L), or teff flour mixed with PBS and incubated at 30 °C for 24 h. A total of 50 µl was transferred into fresh medium (constituting 1% of the final volume) on each of the subsequent five days. The avg ± stdev of three replicate cultures is shown.

To assess the capacity of B1.1 and B1.3 to compete in nutritive conditions resembling their isolation source, the two strains were incubated in an aqueous suspension of teff flour, the primary ingredient in teff flour injera batter. Strains B1.1 and B1.3 grew in the teff flour suspension and reached similar numbers after 24 h incubation (starting at 10^5^ CFU/ml and reaching 4.6 x 10^8^ CFU/ml after 24 h at 30 °C, approximately 4 generations) (data not shown). When co-inoculated into the teff flour, B1.1 and B1.3 were present in equal proportions after 24 h incubation (**Fig. 3D**). However, by the third day of passage, B1.3 reached higher numbers than B1.1, and these differences were sustained for the subsequent days of passage (**Fig. 3D and Fig. S3**). These findings show that despite the smaller genome size and limited biosynthetic capacity of *L. plantarum* B1.3, that strain is highly adapted for growth in nutrient-rich environments and that B1.1 and B1.3 are adapted for co-existence in teff flour.

### *L. plantarum* B1.3 growth in mMRS is not improved with the addition of nucleotides, amino acids, or vitamins

Because the B1.3 genome lacks many biosynthetic pathways and nutrient transporters present in other *L. plantarum* strains and because beef extract and teff flour is replete with different amino acids, carbohydrates, nucleosides, vitamins, and trace metals, it was not possible to predict which nutrient(s) are most important for robust B1.3 growth. Therefore, we performed nutrient addition experiments with mMRS with the goal to identify compounds able to improve growth of B1.3 to levels found in cMRS and comparable to B1.1. These experiments showed that supplementation of exogenous nucleotides (adenine, guanine, and uracil) either singly or combined were not sufficient to increase the growth characteristics of B1.3 in mMRS (**Fig. 4 and Table 2**). Growth was also not improved when amino acids were added to mMRS, either as single amino acid additions (**Fig. S4**) or when provided in a mixture with adenine, inositol, and p-aminobenzoic acid in yeast synthetic drop-out medium supplement (YSDMS) (**Fig. 4 and Table 2**). Moreover, the growth rate of strain B1.3 was not altered in mMRS supplemented with a vitamin mixture (biotin, folic acid, pyridoxine hydrochloride, thiamin HCl, riboflavin, nicotinic acid, D-pantothenate, vitamin B_12_, p-aminobenzoic acid, and thioctic acid) (**Fig. 4 and Table 2**). For *L. plantarum* B1.1, growth of that strain was not affected in mMRS supplemented with nucleotides or amino acids (**Fig. 4 and Table 2 and Fig. S4**). The addition of the vitamin mixture had a negative effect on B1.1, resulting in a reduced growth rate compared to when incubated mMRS (0.53 h^-1^ ± 0.01) (p < 0.05, Student T’s-test) (**Table 2**).

**Fig. 4.**
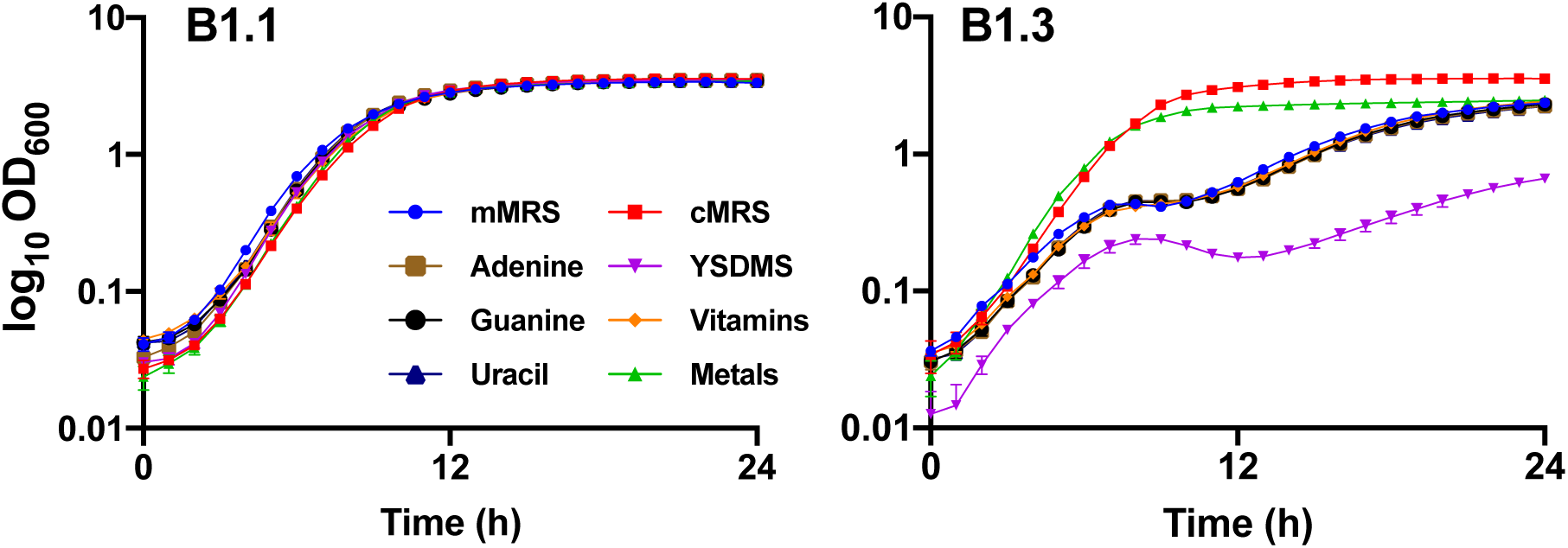
Growth of *L. plantarum* B1.1 and B1.3 in mMRS amended with different nutritional supplements. *L. plantarum* B1.1 and B1.3 were incubated at 30 °C for 24 h in cMRS or mMRS containing additional adenain, guanine, or uracil, yeast synthetic drop-out medium supplement, a mixture of vitamins, or a mixture of trace metals. The avg ± stdev OD_600_ of three replicate cultures is shown.

**Table 2.**
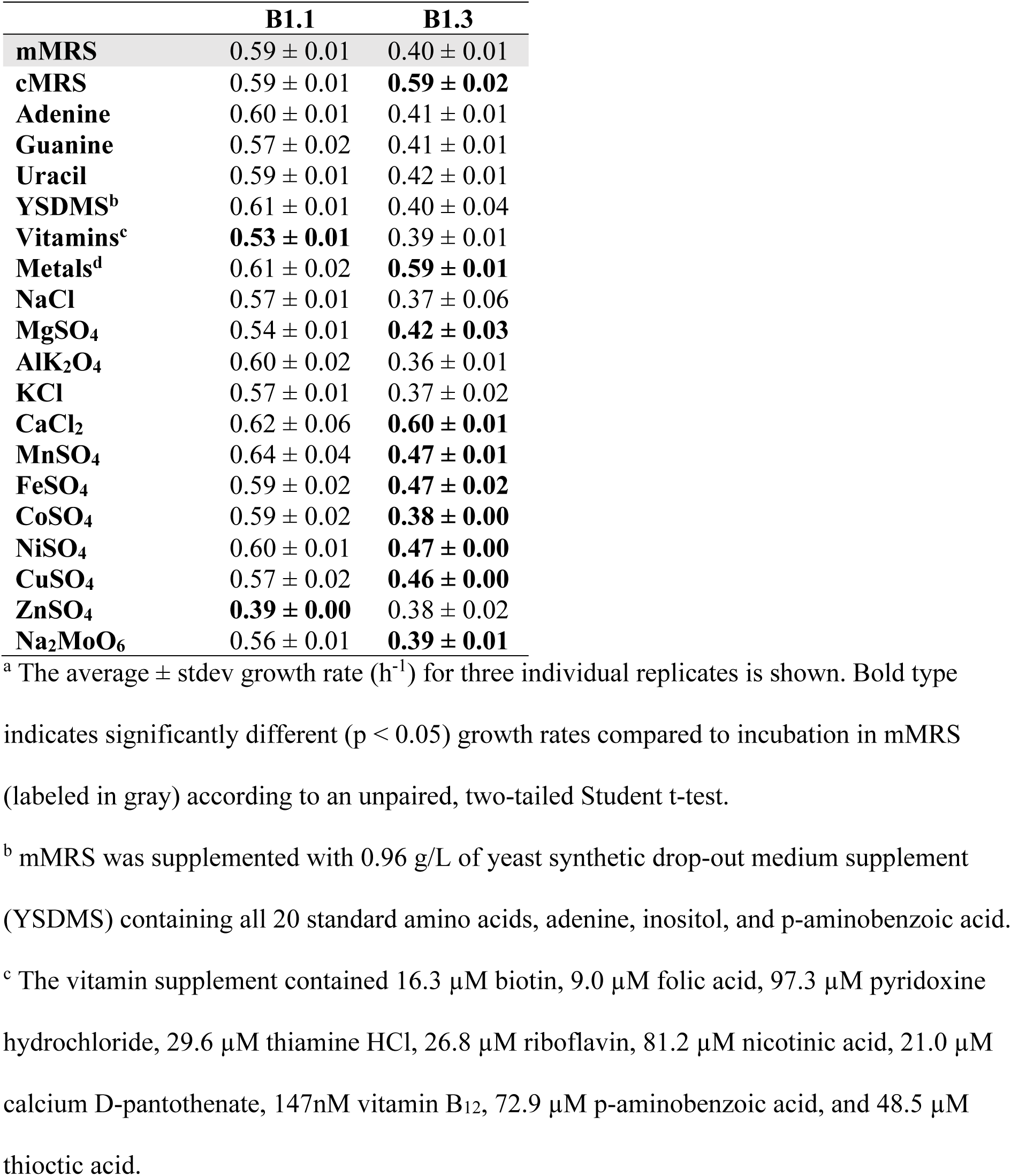
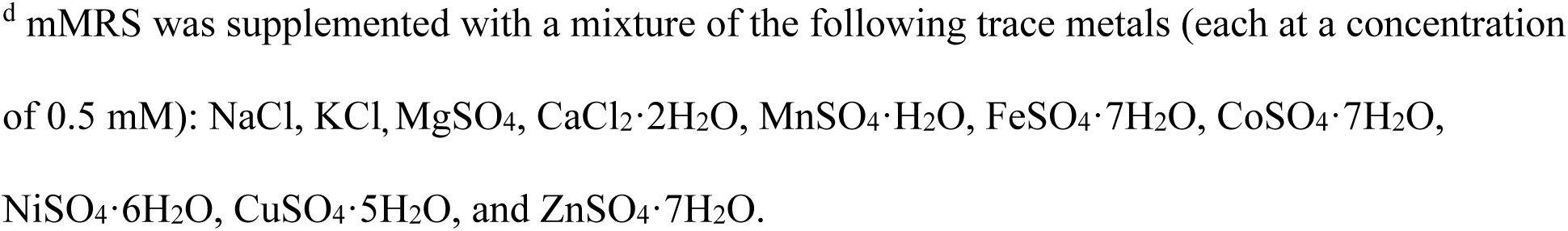
Growth rates of *L. plantarum* B1.1 and B1.3 in MRS supplemented with different nutrients ^a^.

|  | **B1.1** | **B1.3** |
| --- | --- | --- |
| **mMRS** | 0.59 ± 0.01 | 0.40 ± 0.01 |
| **cMRS** | 0.59 ± 0.01 | **0.59 ± 0.02** |
| **Adenine** | 0.60 ± 0.01 | 0.41 ± 0.01 |
| **Guanine** | 0.57 ± 0.02 | 0.41 ± 0.01 |
| **Uracil** | 0.59 ± 0.01 | 0.42 ± 0.01 |
| **YSDMS<sup>b</sup>** | 0.61 ± 0.01 | 0.40 ± 0.04 |
| **Vitamins<sup>c</sup>** | **0.53 ± 0.01** | 0.39 ± 0.01 |
| **Metals<sup>d</sup>** | 0.61 ± 0.02 | **0.59 ± 0.01** |
| **NaCl** | 0.57 ± 0.01 | 0.37 ± 0.06 |
| **MgSO<sub>4</sub>** | 0.54 ± 0.01 | **0.42 ± 0.03** |
| **AlK<sub>2</sub>O<sub>4</sub>** | 0.60 ± 0.02 | 0.36 ± 0.01 |
| **KCl** | 0.57 ± 0.01 | 0.37 ± 0.02 |
| **CaCl<sub>2</sub>** | 0.62 ± 0.06 | **0.60 ± 0.01** |
| **MnSO<sub>4</sub>** | 0.64 ± 0.04 | **0.47 ± 0.01** |
| **FeSO<sub>4</sub>** | 0.59 ± 0.02 | **0.47 ± 0.02** |
| **CoSO<sub>4</sub>** | 0.59 ± 0.02 | **0.38 ± 0.00** |
| **NiSO<sub>4</sub>** | 0.60 ± 0.01 | **0.47 ± 0.00** |
| **CuSO<sub>4</sub>** | 0.57 ± 0.02 | **0.46 ± 0.00** |
| **ZnSO<sub>4</sub>** | **0.39 ± 0.00** | 0.38 ± 0.02 |
| **Na<sub>2</sub>MoO<sub>6</sub>** | 0.56 ± 0.01 | **0.39 ± 0.01** |

<sup>a</sup> The average ± stdev growth rate (h<sup>-1</sup>) for three individual replicates is shown. Bold type indicates significantly different (p < 0.05) growth rates compared to incubation in mMRS (labeled in gray) according to an unpaired, two-tailed Student t-test.
<sup>b</sup> mMRS was supplemented with 0.96 g/L of yeast synthetic drop-out medium supplement (YSDMS) containing all 20 standard amino acids, adenine, inositol, and p-aminobenzoic acid.
<sup>c</sup> The vitamin supplement contained 16.3 µM biotin, 9.0 µM folic acid, 97.3 µM pyridoxine hydrochloride, 29.6 µM thiamine HCl, 26.8 µM riboflavin, 81.2 µM nicotinic acid, 21.0 µM calcium D-pantothenate, 147nM vitamin B<sub>12</sub>, 72.9 µM p-aminobenzoic acid, and 48.5 µM thioctic acid.
<sup>d</sup> mMRS was supplemented with a mixture of the following trace metals (each at a concentration of 0.5 mM): NaCl, KCl, MgSO<sub>4</sub>, CaCl<sub>2</sub>·2H<sub>2</sub>O, MnSO<sub>4</sub>·H<sub>2</sub>O, FeSO<sub>4</sub>·7H<sub>2</sub>O, CoSO<sub>4</sub>·7H<sub>2</sub>O, NiSO<sub>4</sub>·6H<sub>2</sub>O, CuSO<sub>4</sub>·5H<sub>2</sub>O, and ZnSO<sub>4</sub>·7H<sub>2</sub>O.

### Metal salts improve *L. plantarum* B1.3 growth in mMRS

The growth rate of *L. plantarum* B1.3 increased significantly in mMRS containing a mixture of metal salts (0.5 mM of each of the following: NaCl, MgSO_4_, KCl, CaCl_2_, MnSO_4_, FeSO_4_, CoSO_4_, NiSO_4_, CuSO_4_, and ZnSO_4_) (p < 0.05, Student’s T-test) and was equivalent to B1.3 in cMRS (**Table 2 and Fig. 4**). This effect was specific for B1.3 because the growth rate and cell numbers of strain B1.1 were not altered by adding metal salts to the laboratory culture medium (**Table 2 and Fig. 4**).

To assess the contributions of individual metals on *L. plantarum* growth, single metal addition growth experiments were carried out in mMRS. Neither NaCl nor KCl was sufficient to improve the growth characteristics of B1.3 at either 5 mM (**Table 2 and Fig. 5**) or at higher concentrations (50 mM or 100 mM) (data not shown). Although the presence of supplemental 5 mM AlKS_2_O_4_ resulted in significantly higher B1.3 final cell numbers (indicated by the OD_600_ after 24 h incubation; p < 0.05, Student’s T-test) (**Fig. 5**), the growth rate of that strain in mMRS with the aluminum salt was not changed compared to mMRS (**Table 2**). Instead, the growth rates and final OD_600_ values of B1.3 were significantly higher during incubation in mMRS containing either additional 5 mM MgSO_4_, CaCl_2_, MnSO_4_, FeSO_4_, CoSO_4_, NiSO_4_, CuSO_4_, ZnSO_4_, or Na_2_MoO_6_ (p < 0.05, Student’s T-test) (**Table 2 and Fig. 5**). Inclusion of those divalent cation metal salts also eliminated the diauxic growth of B1.3 in mMRS (**Fig. 5**).

**Fig. 5.**
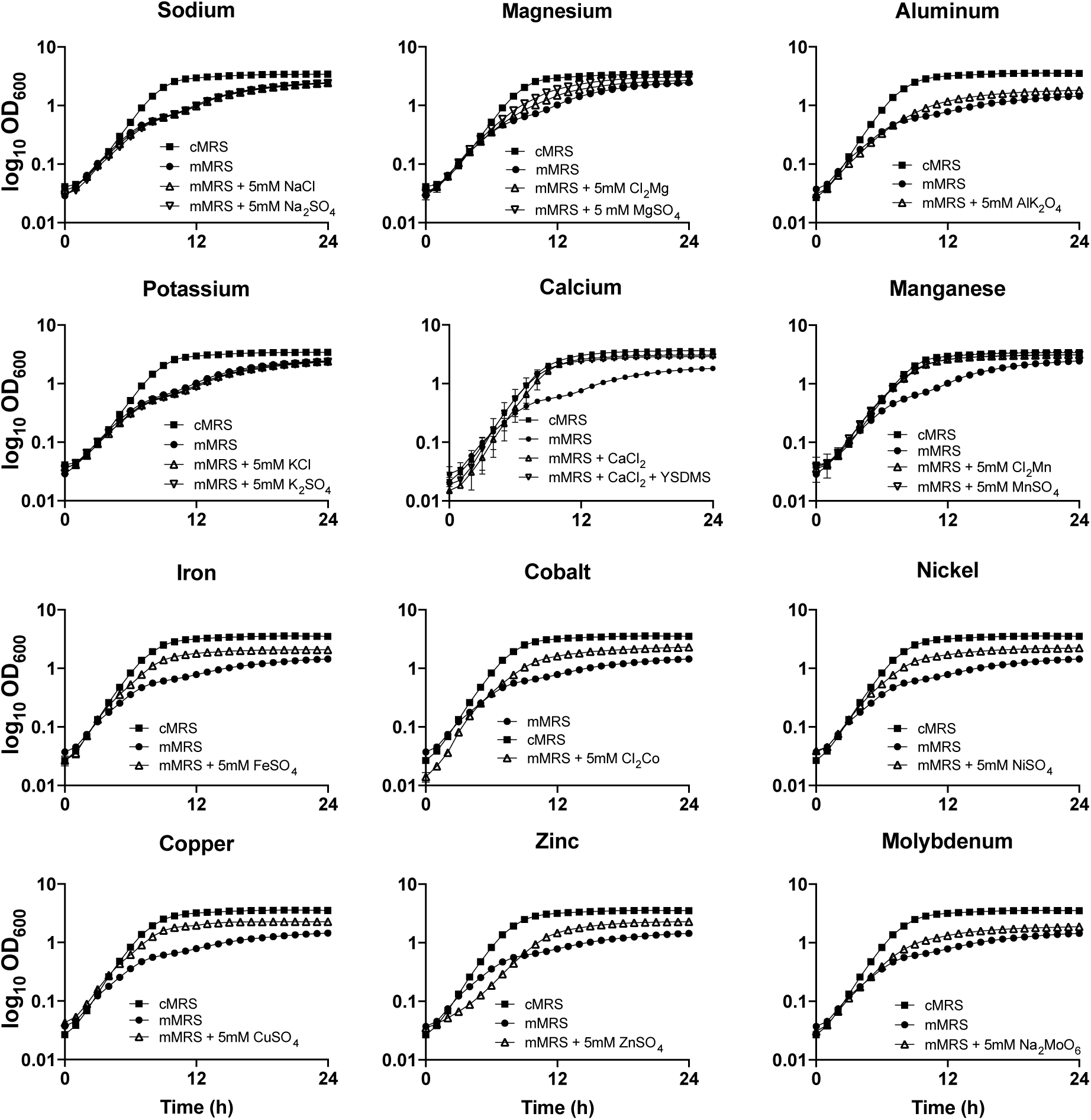
Growth of *L. plantarum* B1.3 in mMRS supplemented trace metals. *L. plantarum* B1.3 were inoculated into mMRS containing 5mM of individual trace metals and incubated at 30 °C for 24 h. The avg ± stdev OD_600_ of three replicate cultures is shown.

Among the metal salt amendments, 5 mM CaCl_2_ resulted in the greatest improvement in B1.3 growth in mMRS. Surprisingly, this single modification improved the growth rate of B1.3 in mMRS (0.60 h^-1^ ± 0.01) to the extent that it was equal to the growth rate of this strain in cMRS (0.62 h^-1^ ± 0.01) and MRS-BE (0.61 h^-1^ ± 0.01) (**Table 2**). This change occurred without increasing final cell numbers (final OD_600_ in mMRS-Ca (3.06 ± 0.03) compared to cMRS (3.60 ± 0.03) or mMRS-BE (3.36 ± 0.05)). CaCl_2_ associated improvements in the growth rate of B1.3 were not due to increased concentrations of the chloride or sulfate anions in mMRS, as shown by their lack of effect in experiments with NaCl and KCl and by the equivalent growth of B1.3 in mMRS containing the Mg^2+^ and Mn^2+^ salts of both of those anions (**Fig. 5**). For *L. plantarum* B1.1, the addition of the individual metal salts in mMRS either did not change the growth rate or had a negative impact (5 mM ZnSO_4_) on the growth characteristics of that strain (**Table 2 and Fig. S5**).

### Intracellular metal concentrations are reduced in *L. plantarum* B1.3

To better understand why growth of strain B1.3 in mMRS was improved by the addition of divalent metal cations, intracellular metal concentrations for early stationary-phase B1.1 and B1.3 cells grown in cMRS and mMRS were quantified by inductive coupled plasma mass spectrometry (ICP-MS). Intracellular sodium, magnesium, aluminum, potassium, calcium, manganese, iron, copper, and zinc metal concentrations were detectable above background levels by ICP-MS. Cobalt, molybdenum, and nickel were below the detection limit (data not shown). ICP-MS showed that in cMRS the sum of those metal concentrations was equivalent between B1.1 and B1.3 (**Fig. 6**). In mMRS, B1.1 contained 2.7-fold higher levels than B1.3 (**Fig. 6**).

**Fig. 6.**
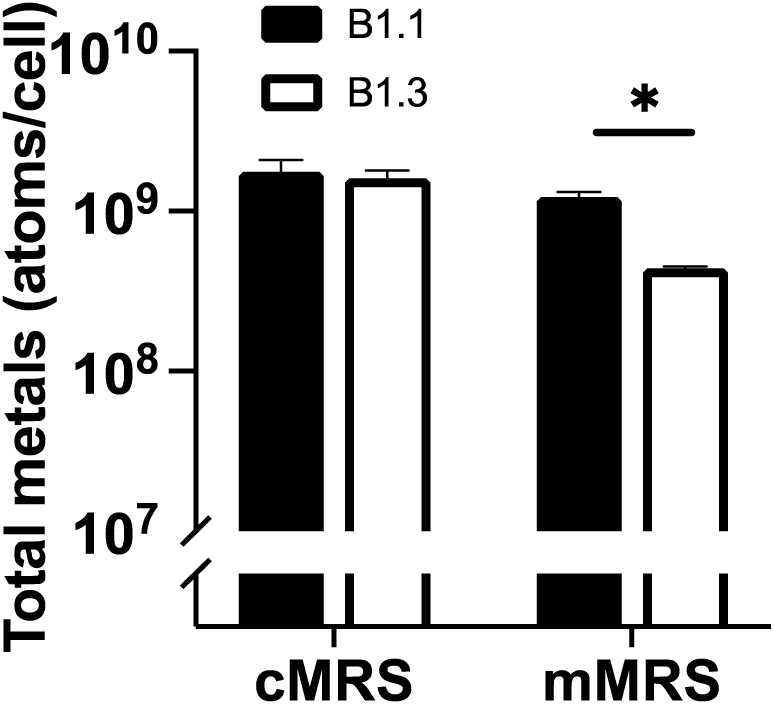
Sum of intracellular metal concentrations in *L. plantarum* B1.1 and B1.3 in cMRS or mMRS as determined by ICP-MS. Intracellular metal concentrations after growth in cMRS (black bars) and mMRS (open bars) are shown. Total metal concentrations were calculated by adding the concentrations of sodium, magnesium, aluminum, potassium, calcium, manganese, iron, cooper, and zinc atoms. Levels of cobalt and nickel, and molybdenum were below the detection limit for ICP-MS. The avg ± stdev of three replicate cultures is shown. Significant differences were calculated using a Student’s T-test (p < 0.05).

Sodium and potassium were the most abundant metals in *L. plantarum* B1.1 and B1.3 in both mMRS and cMRS (between 5 x 10^8^ to 1 x 10^9^ atoms / cell) (**Fig. 7**). Within the range of detection, copper and iron were the least abundant (between 1 x 10^4^ to 7 x 10^4^ atoms / cell) (**Fig. 7**). MRS growth medium type did not affect the intracellular metal concentrations of strain B.1.1 (**Fig. 7**). As expected, based on total intracellular metal concentrations (**Fig. 6**), B1.3 contained significantly lower quantities of sodium, magnesium, potassium, calcium, manganese, copper, and zinc when grown in mMRS relative to cMRS (**Fig. 7**) (p < 0.05 Student’s T-test). However, aluminum was increased (**Fig. 7**).

**Fig. 7.**
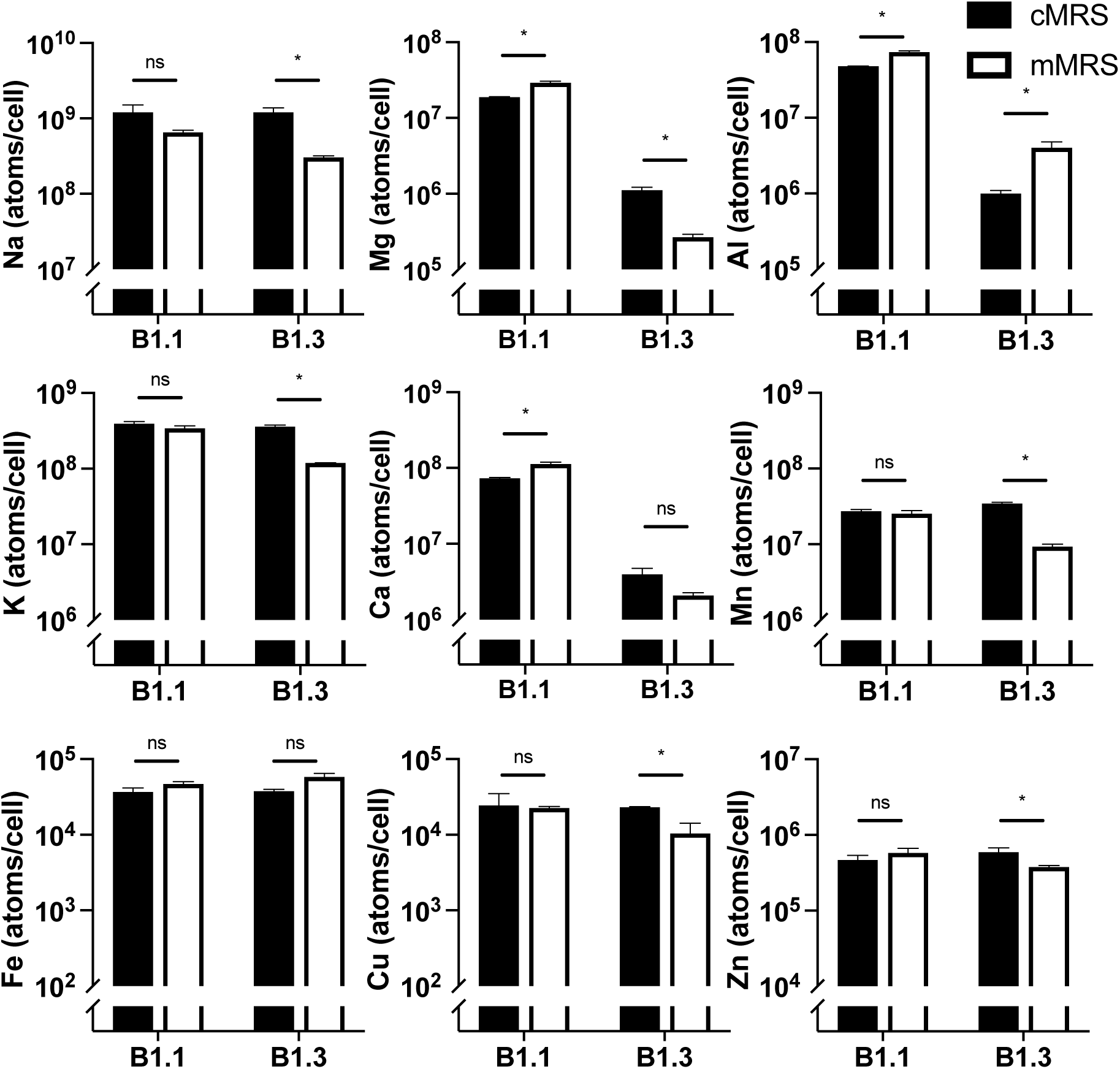
Intracellular metal concentrations in *L. plantarum* B1.1 and B1.3 in cMRS or mMRS as determined by ICP-MS. Intracellular metal concentrations of B1.1 (black bars) and B1.3 (open bars) after 24 h of growth in cMRS and mMRS are shown as determined by ICP-MS. The avg ± stdev of three replicate cultures is shown. Significant differences were calculated using a Student’s T-test (p < 0.05).

Comparisons of intracellular metal concentrations between the *L. plantarum* B1.1 and B1.3 strains showed that in cMRS *L. plantarum* B1.3 contained less aluminum (48-fold), calcium (18-fold), magnesium (17-fold), and higher amounts of manganese (1.5-fold) than strain B1.1 (**Fig. 7**). In mMRS, the quantities of eight out of the nine individual metals were present in significantly lower quantities in B1.3 relative to B1.1, with magnesium (110-fold), calcium (55-fold), and aluminum (18-fold) and constituting the greatest difference from B1.1 (**Fig. 7**). These results show that *L. plantarum* B1.3 and B1.1 have different capacities to transport and retain metals in the cell and intracellular metal concentrations are culture media dependent.

### *L. plantarum* B1.3 exhibits robust growth and competitive fitness in mMRS amended with supplemental calcium

The robust growth of *L. plantarum* B1.3 in mMRS containing supplemental 5mM CaCl_2_ and the low quantities of calcium detected in that strain in cMRS and mMRS led us to investigate the specificity for this calcium requirement. First, we found that the inclusion of 5 mM CaCl_2_ was sufficient to increase the growth rate and cell yield of strain B1.3 in mMRS. Growth was not further improved when YSDMS was added to the laboratory culture medium (**Fig. 4**). Moreover, the growth rate of B1.3 in mMRS containing 5mM CaCl_2_ was impaired when EGTA (ethylene glycol-bis (β-aminoethyl ether)-N,N,N′,N′-tetraacetic acid), a metal chelator with a high affinity for Ca^2+^ was included. The growth rates of *L. plantarum* B1.1 and B1.3 were reduced in mMRS EGTA concentration-dependent manner when either 5 mM, 10 mM, or 25 mM EGTA was added (**Fig. S6**). The negative effect of EGTA was prevented (B1.1) or minimized (B1.3) when CaCl_2_ was also included in the mMRS (**Fig. 8**). Additionally, stationary phase B1.3 did not flocculate and instead remained suspended in the CaCl_2_ amended mMRS. Changes in flocculation occurred in a CaCl_2_ concentration-dependent manner, such that no aggregation was observed in the presence of 2.5 mM CaCl_2,_ and intermediate aggregation phenotype was found for 1 mM CaCl_2_, and the cells aggregated at lower calcium (0.5 mM CaCl_2_) concentrations (**Fig. S7**). This effect was specific for calcium, because B1.3 cell aggregates formed when that strain was grown in mMRS containing MgSO_4,_ AlK_2_O_4_, or NaCl (**Fig. S8**). By comparison, *L. plantarum* B1.1 remained suspended in all mMRS conditions (**Fig. S9**).

**Fig. 8.**
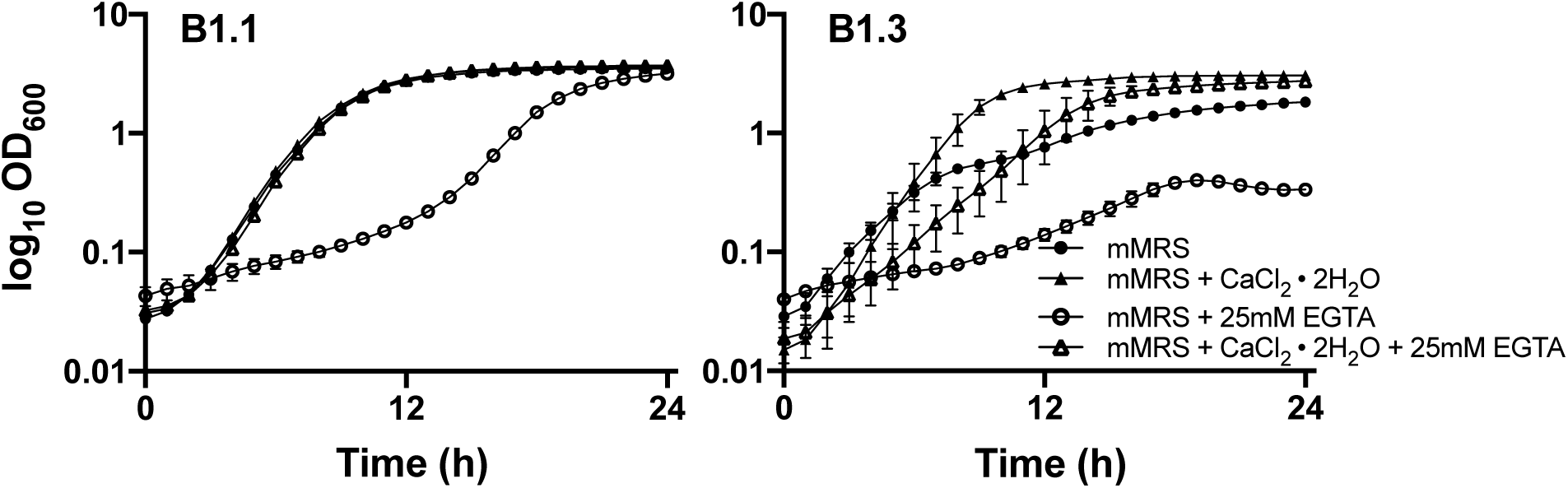
Growth of *L. plantarum* B1.3 is significantly impacted by the addition of EGTA, a metal chelator, but addition of calcium chloride ameliorates that affect. *L. plantarum* B1.1 and B1.3 were inoculated into mMRS, mMRS supplemented with 5 mM CaCl_2_ꞏ2H_2_O with or without 25 mM EGTA and were incubated at 30 °C for 24 h. The avg ± stdev OD_600_ of three replicate wells are shown.

Lastly, we examined the effect of calcium on the competitive fitness of B1.1 and B1.3 in co-culture in mMRS supplemented with 5 mM CaCl_2_. Consistent with the improved growth of B1.3 in that laboratory culture medium, cell numbers of B1.3 and B1.1 were sustained in equal ratios (**Fig. 9**). The numbers of B1.3 were also significantly higher than B1.1 after three days of successive passage (p < 0.05 Student’s T-test). In mMRS containing either 5mM NaCl, MgSO_4_, or AlK_2_O_4_, B1.3 was outcompeted by B1.1 (**Fig. S10**). Our findings therefore confirm that sustaining both B1.3 and B1.1 in co-culture is dependent on calcium availability.

**Fig. 9.**
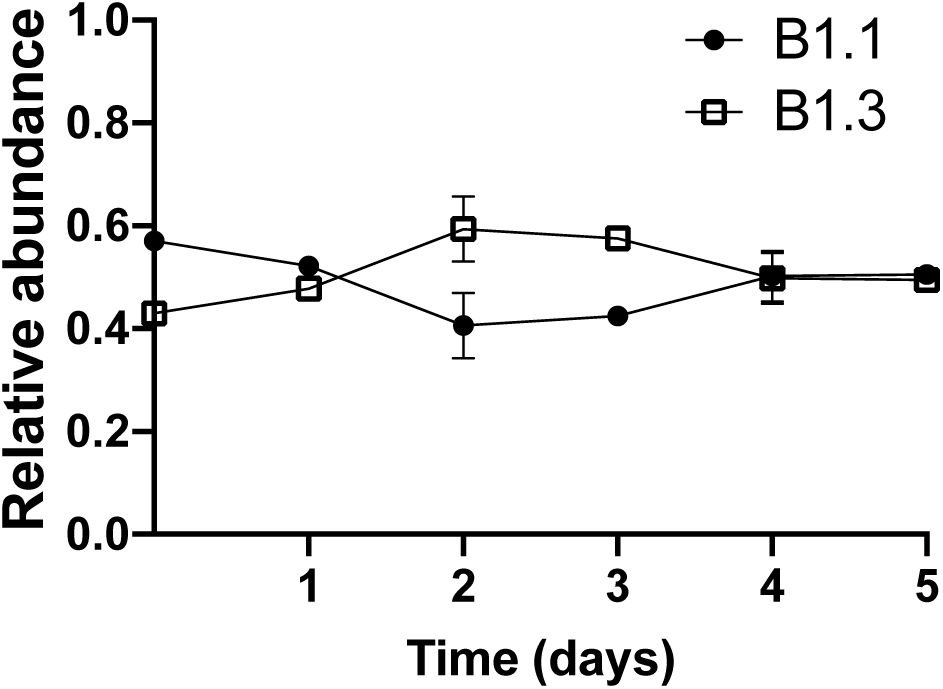
Competitive growth f *L. plantarum* B1.1 and B1.3 in mMRS amended with 5mM CaCl_2_ꞏ2H_2_O. Equal numbers of *L. plantarum* B1.1 and B1.3 (10^5^ CFU/ml) were co-inoculated in mMRS supplemented with 5mM CaCl_2_ꞏ2H_2_O incubated at 30 °C for 24 h. A total of 50 µl was transferred into fresh medium (constituting 1% of the final volume) on each of the subsequent five days. The avg ± stdev of three replicate cultures are shown.

## Discussion

Food fermentations are complex microbial ecosystems which frequently contain multiple, highly related species and strains under constant selective pressure due to the changing physicochemical conditions in the food matrix. The co-occurrence of highly-related phylogenetic clusters is not well understood, but presumably is affected by both biotic and abiotic niche partitioning as well as mutualistic and antagonistic interactions (23, 24). These relationships are dependent on the prevailing nutrients in the food environment. In this study, we characterized *L. plantarum* strains B1.1 and B1.3 isolated from teff flour injera batter. These strains are genetically distinct and, due to its smaller size and large mobilome, strain B1.3 appears to have undergone genome reduction, potentially as a consequence of adaptation to teff plants, flour, or injera fermentation batter. However, despite the growth impairments of B1.3 in mMRS, B1.3 was sustained in higher numbers than the more robust strain B1.1 in teff flour and outcompeted B1.1 by a ratio of three to one in cMRS. By examining the growth requirements of those strains, we identified the importance of calcium for competitive fitness of *L. plantarum* B1.3.

Because *L. plantarum* B1.1 and B1.3 were randomly selected colony isolates from a single sample of teff injera batter, we expected that they would exhibit similar genotypes and growth characteristics in laboratory culture medium. Instead, these strains spanned the range of known features of the *L. plantarum* species. Whereas the genome content of B1.1 is similar to the model strain *L. plantarum* WCFS1 and other plant-associated LAB (20), B1.3 contains a highly reduced genome lacking genes encoding for the synthesis of amino acids and vitamins and the metabolism of a variety of carbohydrates. Like other LAB, genome rearrangements in B1.3 appear to have been the result of the acquisition of transposases and other mobile element-mediated gene disruptions (5). The limited biosynthetic capacity of B1.3 is consistent with the evolution of the *Lactobacillales* towards gene loss and metabolic simplification (5, 8, 13, 15–17). The selective pressures in teff flour may be particularly great for genome reductions, because although foods are typically rich sources of nutrients for microbial growth, teff flour has higher concentrations of amino acids and vitamins compared to other cereal grains (barley, wheat, rye, maize, brown rice, sorghum, and pearl millet) (25).

*L. plantarum* B1.3 also grew poorly in mMRS, a modified version of MRS lacking beef extract. Because beef extract contains undefined concentrations of mono- and disaccharides which support *L. plantarum* growth, we previously measured carbohydrate preferences and stress tolerance characteristics of B1.3, B1.1 and other plant-associated *L. plantarum* strains, using mMRS as our reference medium (20). Despite the limitations of using mMRS for assessments of Bl.3, differences between B1.3 and B1.1 in mMRS and cMRS provided an opportunity to identify the nutrients upon which B1.3 is particularly reliant. Remarkably, the provision of additional amino acids, nucleotides, and vitamins did not improve the growth of B1.3 in mMRS. The lack of effect in that medium, may be due to the fact that mMRS is nutrient-replete containing undefined concentrations of those compounds in yeast extract and enzymatic digests of animal proteins (26).

Only with the addition of divalent metal salts to mMRS, either singly or combined, was there an improvement in *L. plantarum* B1.3 growth. The growth rates of B1.3 in mMRS with these elevated metal concentrations increased to levels similar to those found in cMRS, cMRS-BE, and for B1.1. The differences in cation metal requirements between B1.3 and B1.1 were similarly confirmed by ICP-MS, showing that B1.3 cells contain significantly lower intracellular metal concentrations in mMRS. The altered intracellular metal concentrations between the two strains were not limited to the mMRS culture medium. Despite having similar total quantities of metals in cMRS, B1.3 still contained lower amounts of magnesium, aluminum, and calcium compared to strain B1.1. Presently, aside from a few reports suggesting different metal requirements for lactobacilli (27–31), the importance of metals for LAB growth and ecological interactions are not well understood. The requirement of *L. plantarum* for manganese has been specifically investigated (30, 32), likely because *L. plantarum* has a high requirement for Mn^2+^ compared to other bacteria and Mn^2+^ is known to confer resistance against oxidative (33, 34) and heat (35) stress. However, the potentially overlapping effects of metal cations (36) on *L. plantarum* and other lactobacilli remain to be determined. Although *L. plantarum* genomes contain a variety of metal transporters as well as enzymes and processes with requirements for specific metals (21, 22), these pathways are not sufficiently elucidated to identify the specific genes required for metal homeostasis in the *L. plantarum* strains examined here.

Surprisingly, supplementing mMRS with CaCl_2_ resulted in the highest growth rate of B1.3 and enabled that strain to outcompete strain B1.1. Calcium is essential for intracellular signaling and regulation of multiple cellular processes including cell division and development, motility, stress response, and host-pathogen interactions (37, 38). Because the addition of calcium in mMRS prevented the flocculation of *L. plantarum* B1.3 during stationary phase, the requirement for calcium may have been due to calcium-dependent effects on cell surface composition. To this regard, calcium and magnesium are required for bacterial cell wall and teichoic acid stability (39). Calcium was also shown to be important for aggregation, biofilm formation, and adhesion in both Gram negative (e.g. *Xylella fastidiosa* and *Vibrio*) (40–42) and Gram positive bacteria (e.g. *Bacillus subtilis* and *Mycobacterium smegmatis*) (39, 41, 43, 44). In yeast, calcium regulates a flocculation phenotype in a lectin-specific manner (45). Although the specific role of calcium for B1.3 and B1.1 in teff flour fermentations remains to be determined, teff grains also have approximately three-fold higher levels of calcium than sorghum, the cereal grain with the next highest quantities of calcium (50mg/100g) (25). This nutritional requirement of B1.3 for calcium therefore provides further evidence of the adaptation of this strain to teff-associated habitats. Moreover, because both strains were inhibited when EGTA, a chelator with a high affinity for Ca^2+^, was included in the medium, these findings also indicate a broader role for calcium in *L. plantarum,* besides the strain-specific effects observed here.

In conclusion, by examining the nutritional requirements of B1.3, we identified the significance of divalent metal ions and particularly Ca^2+^ on the ecological fitness of *L. plantarum*. Because metals cannot be synthesized or degraded and may damage the cell if present in high quantities (46, 47), tight control over metal homeostasis is likely required by this species as well as other LAB. The importance of trace metals on the growth and auto-aggregation characteristics likely has a role in nutrient-rich food fermentations. These findings improve our understanding of the coexistence of multiple strains of the same microbial species within a single ecosystem as well as emphasize the importance of environmental context when evaluating inter- and intra-species diversity and functionality of LAB in food fermentations.

## Materials and Methods

### Bacterial strains and growth conditions

*L. plantarum* strains B1.1 and B1.3 were previously isolated from Ethiopian teff injera (Yu *et al*., 2021) and were maintained as frozen glycerol stocks at -80 °C. *L. plantarum* was routinely grown at 30 °C in a commercial preparation of MRS (cMRS) (BD, Franklin, NJ), a modified MRS (de Man *et al.*, 1960) (mMRS) was prepared as lacking beef extract, mMRS supplemented with beef extract (8 g/L) (mMRS-BE), or a teff flour medium. To prepare the teff flour medium, whole grain teff flour (Bob’s Red Mill, Milwaukie, Oregon) was sterilized by autoclaving for one hour and then drying for approximately 15 h at room temperature. Teff flour medium was prepared by mixing the teff flour with phosphate buffered saline (PBS, 137 mM NaCl, 2.7 mM KCl, 4.3 mM Na_2_HPO_4_-7H_2_O, 1.4 mM KH_2_PO_4_) (pH 7.2) at a dough yield (100 * weight [flour+PBS]/weight [flour]) (48) of 300.

### Genome comparisons of B1.3

Assembled and annotated genome sequences of *L. plantarum* B1.1 (WWCZ00000000) and B1.3 (WWCY00000000) were retrieved from National Center for Biotechnology Information (https://www.ncbi.nlm.nih.gov/). Comparative genomics analysis using PATRIC was utilized to identified differences in the presence and absence of protein families as compared to seven strains of plant-associated *L. plantarum* (1B1, K4, 8.1, AJ11, BGM37, EL11, WS1.1) (Yu *et al*., 2021) and model strain WCFS1 (21) (https://www.patricbrc.org). Functional genomics analysis was preformed using ANVI’O 6.1 to cluster genes with similar function and categories them into different functional categories (49). Genome size comparison were made against the 663 genome assemblies available on National Center for Biotechnology Information database in January 2022 (https://www.ncbi.nlm.nih.gov/).

### Competitive fitness of B1.1 and B1.3 between *L. plantarum* in different laboratory media and teff flour

*L.* plantarum B1.1 and B1.3 were first incubated in cMRS for 24 h at 30 °C. The cells were then collected by centrifugation at 5,000 x g for 5 min, washed twice in PBS (pH 7.2), and then suspended in cMRS, mMRS, mMRS supplemented with 5mM CaCl_2_ꞏ2H_2_O, or teff flour medium at a starting density of 10^5^ CFU/mL and grown for 24 h at 30°C. Cultures were then propagated at 1% into fresh media every 24 h to simulate back-slopping. Serial dilutions of cultures after 24 h were plated on mMRS containing 2% (w/v) (58mM) sucrose (mMRS-sucrose) and incubated at 48 h at 30 °C. B1.1 and B1.3 colony size and shape on mMRS-sucrose were used to identify the two strains.

### *L. plantarum* growth requirements in mMRS

*L. plantarum* B1.1 and B1.3 were incubated in cMRS for 24 h at 30 °C, collected by centrifugation at 5,000 x g for 5 min, washed twice in PBS (pH 7.2), and then suspended in mMRS. The cell suspensions were then distributed into 96-well microtiter plates (Thermo Fisher Scientific, Waltham, MA) at an optical density (OD) at 600 nm (OD_600_) of 0.02. Wells contained mMRS supplemented with exogenous nucleobases (adenine, guanine, or uracil), vitamins, amino acids, or metals. Controls included mMRS diluted with an equal volume of water (5% (v/v)) instead of nutritional supplements. Growth was measured using a Synergy 2 microplate reader (Biotek, Winooski, VT) at 30 °C for 36 h every hour.

For experiments with the nucleobases, mMRS was amended with purines (adenine (148 µM) and guanine (132µM)) or the pyrimidine (uracil (178µM). For incubation in a supplemental vitamin mixture, mMRS was amended to a final concentration of 16.3 µM biotin, 9.0 µM folic acid, 97.3 µM pyridoxine hydrochloride, 29.6 µM thiamine HCl, 26.8 µM riboflavin, 81.2 µM nicotinic acid, 21.0 µM calcium D-pantothenate, 147nM vitamin B_12_, 72.9 µM p-aminobenzoic acid, and 48.5 µM thioctic acid.

For assessments of growth with supplemental amino acids, mMRS was amended with yeast synthetic drop-out medium supplement (YSDMS) (0.96 g/L) (Sigma-Aldrich, St. Louis, MO) or Casamino acids (BD, Franklin, NJ) (15mg/mL). YSDMS contains all 20 amino acids, adenine, inositol, and p-aminobenzoic acid. For the single addition amino acids, the concentrations of amino acids provided were at five-times the concentration was described previously (18), with the exception of tyrosine, which was supplemented at the previously described concentrations (Teusink *et al.*, 2005) to its limited solubility in the growth medium.

To assess the effects of added metal salts, the following metal salts were used: CaCl_2_ꞏ2H_2_O, K_2_SO_4_ MgSO_4,_ MnSO_4_ꞏH_2_O, NaCl, Na_2_MoO_6,_ NiSO_4_ꞏ6H_2_O, and ZnSO_4_ꞏ7H_2_O (Fisher Scientific, Pittsburg, PA); AlK_2_O_4,_ Cl_2_Mgꞏ6H_2_O, Cl_2_Mnꞏ4H_2_O, KCl, and Na_2_SO_4_ (Sigma-Aldrich, St. Louis, MO), FeSO_4_ꞏ7H_2_O (Spectrum, New Brunswick, NJ), CoSO_4_ꞏ7H_2_O (Acros Organics, Geel, Belgium), and CuSO_4_ꞏ5H_2_O (VWR International, Radnor PA).

### ICP-MS

*L. plantarum* B1.1 and B1.3 were incubated in cMRS and mMRS for 24 h at 30 °C. A total of 3.16 x 10^9^ cells were collected by centrifugation at 5,000 x g for 5 min and washed twice in PBS. The resulting cell materials was digested by incubating at 95 °C for 45 minutes in a 60% concentrated trace metal grade HNO_3_, allowed to cool, then diluted with MilliQ water to a final concentration of 6% HNO_3,_ and analyzed with an Agilent 7500Ce ICP-MS (Agilent Technologies, Palo Alta, CA) for simultaneous determination of select metals (Na, Mg, Al, K, Ca, Mn, Fe, Co, Ni, Cu, Zn) at the UC Davis Interdisciplinary Center for Plasma Mass Spectrometry (http://icpms.ucdavis.edu/). Raw counts (ppb, ng/ml) of each select metal were converted to the number of atoms using the appropriate element molecular weights and normalized by CFU to obtain atoms per cell.

### Auto-aggregation, flocculation assay

*L. plantarum* B1.1 and B1.3 were incubated in 5mL of cMRS, mMRS, or mMRS supplemented with 5mM CaCl_2_ꞏ2H_2_O, 5mM NaCl, 5mM MnSO_4_ꞏH_2_O, or 5mM AlK_2_O_4_ (in triplicate) at 30°C for 24h. The cultures were then vortexed vigorously for 10 s and the OD_600_ was measured by collecting cells with a pipette tip placed 2 cm below the surface of the laboratory culture medium. The cultures were then incubated at 30°C and the OD_600_ was measured again 24 and 48h later. The percentage of aggregation was calculated using the formula, [1 - (Final OD_600_/Overnight OD_600_)] x 100, where Final OD_600_ represents the optical density 2 cm below the surface at the 48 h or 72 h and Overnight OD_600_ represents the optical density 2 cm below the surface at 24 h from initial inoculation. At the end of the incubation period, the cultures were imaged to visualize the differences in the auto-aggregation, flocculation phenotype.

## Acknowledgements

We are grateful to the USDA National Institute of Food and Agriculture (Grant No. 2015-67017-23116) for funding this work. We would like to thank Menkir Tamrat and Monica Spiller for providing the injera used to isolate the *L. plantarum* strains.

---

**Table S1.**
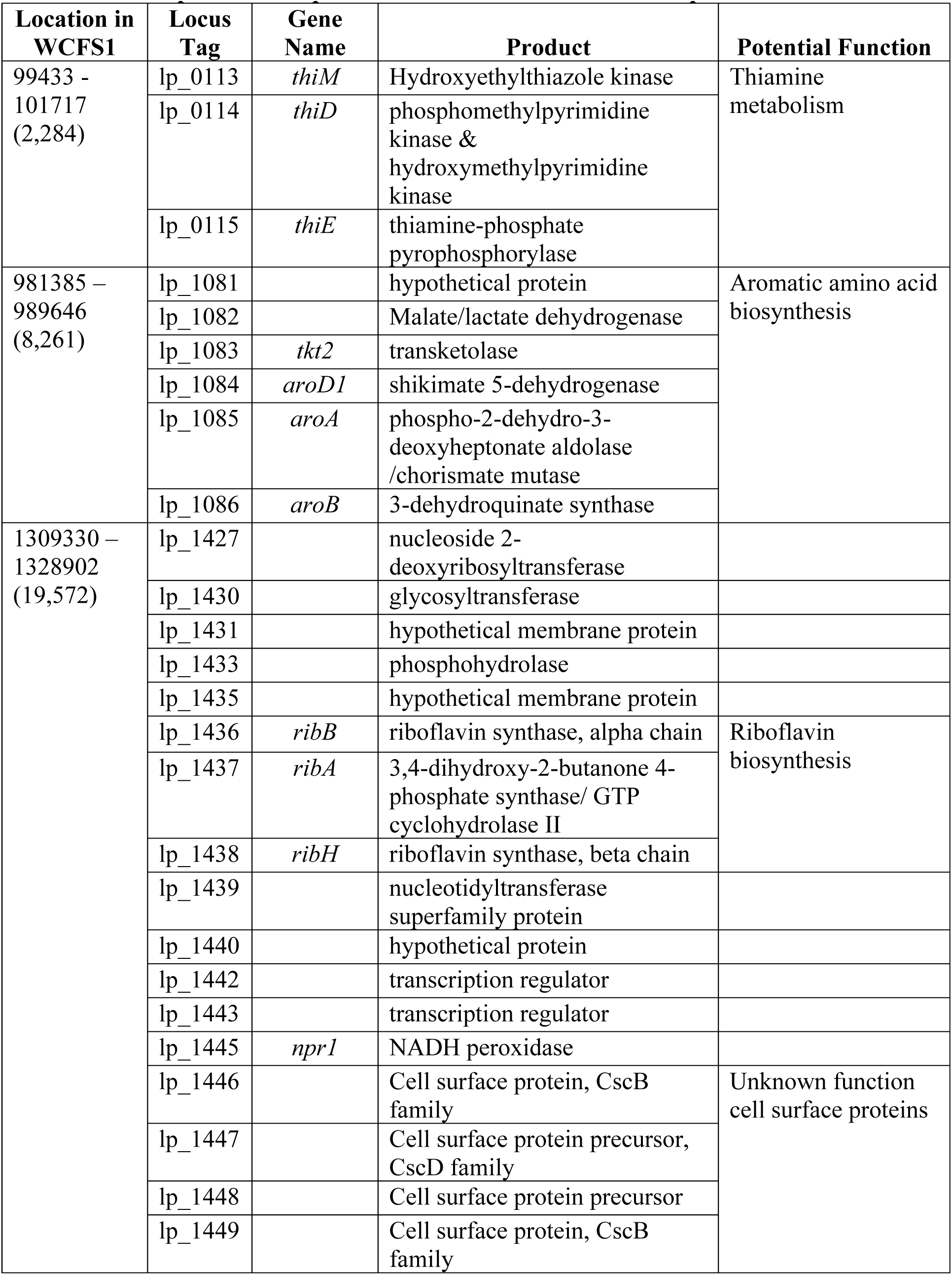
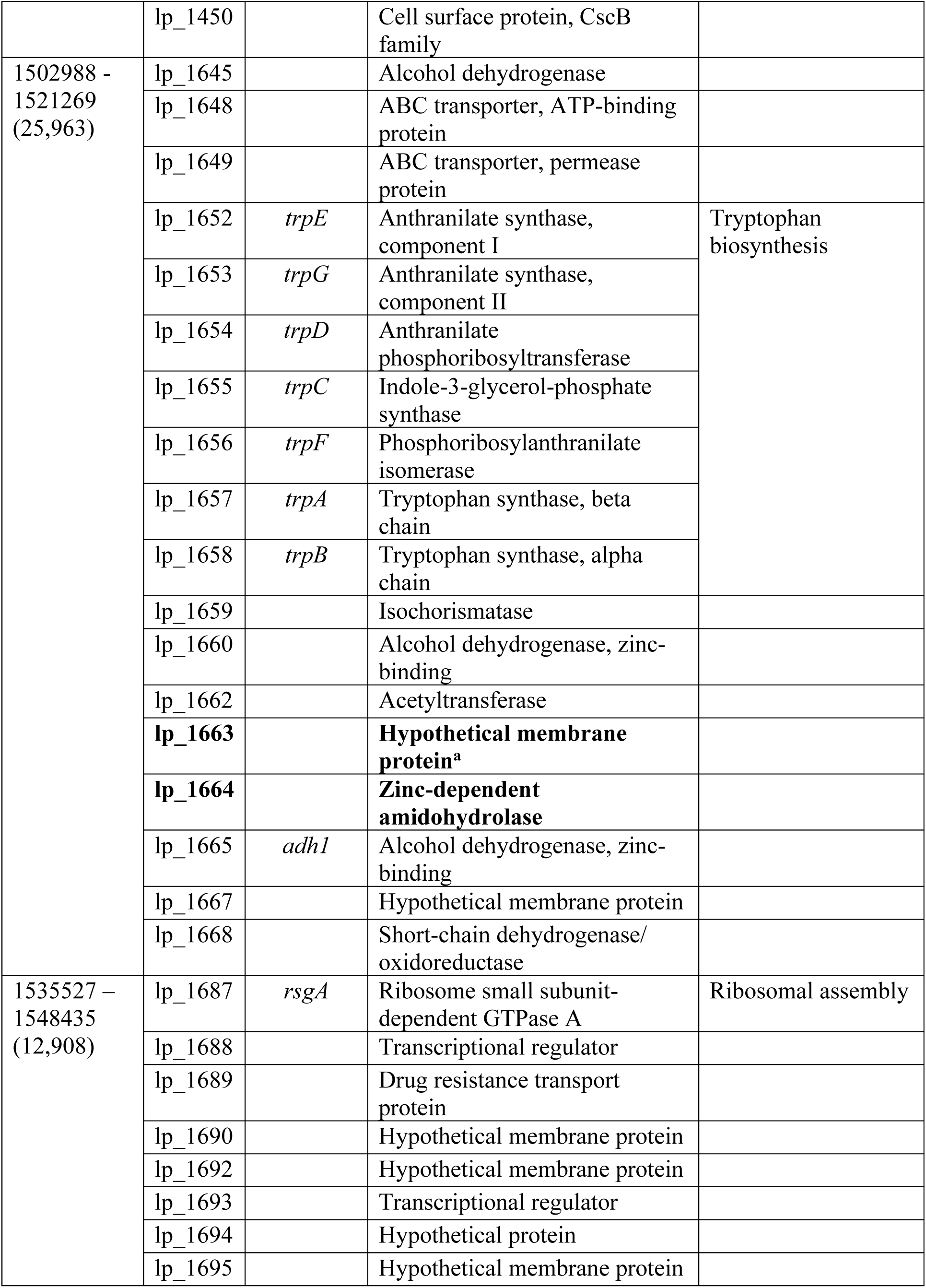
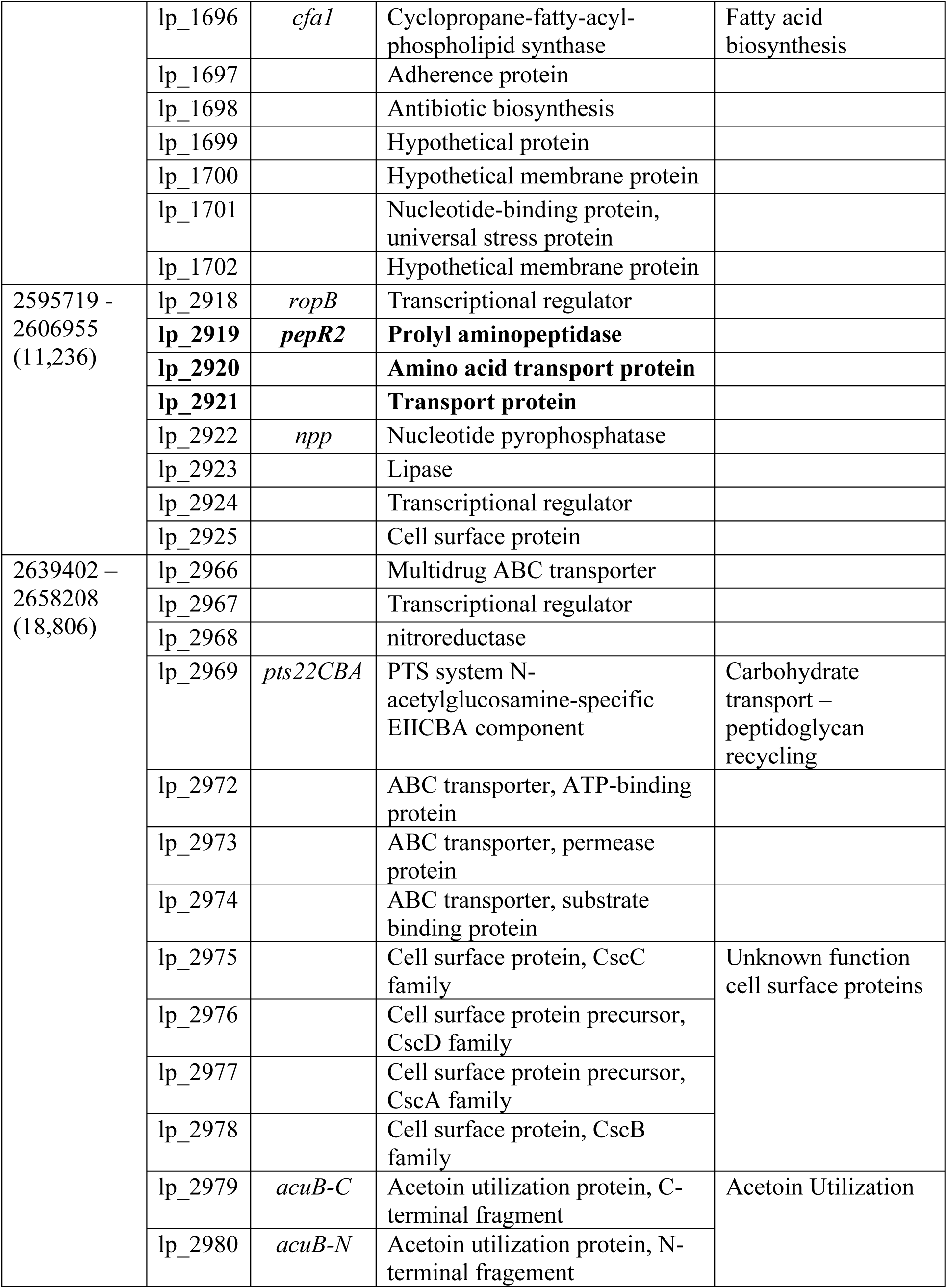
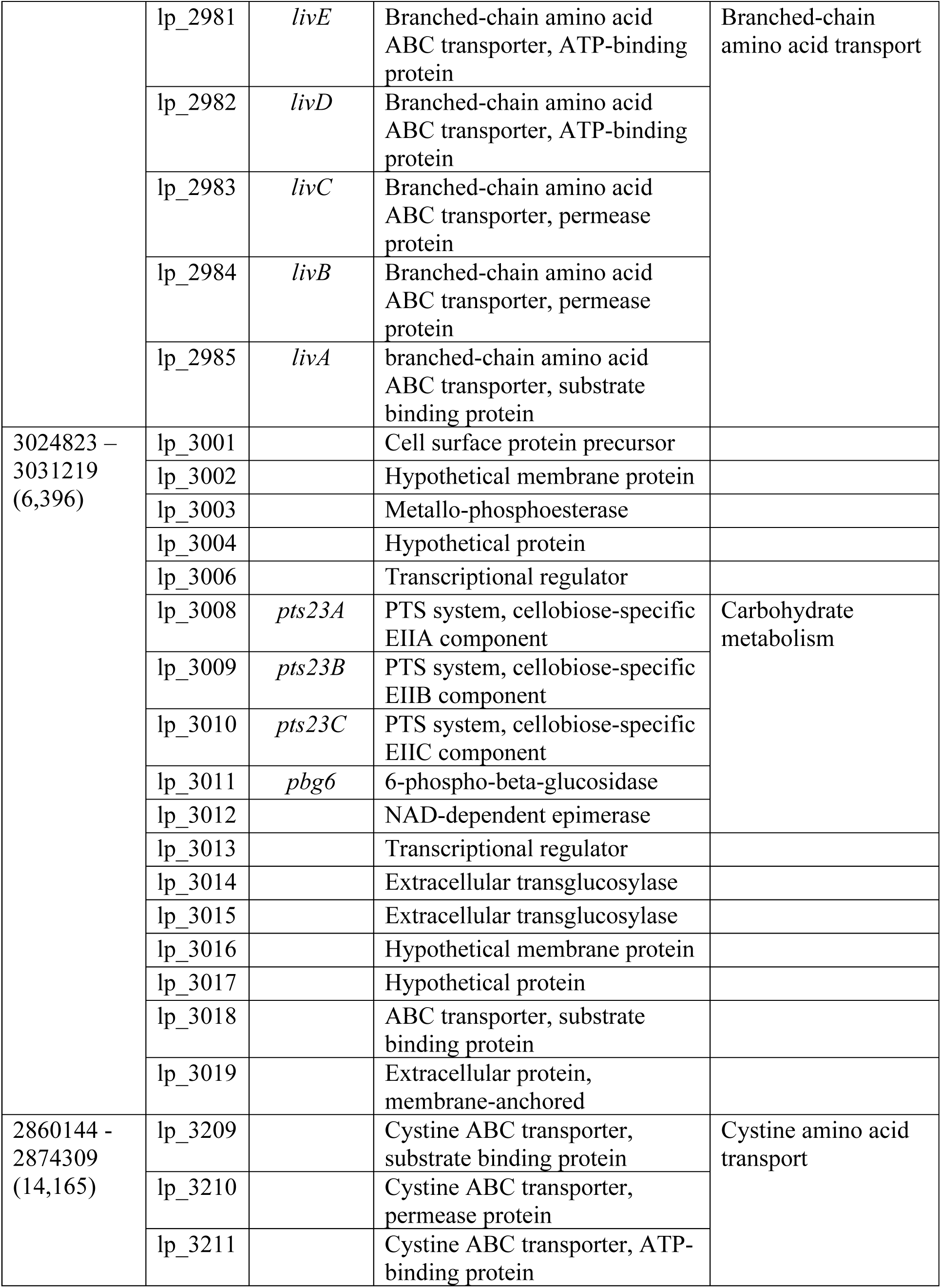
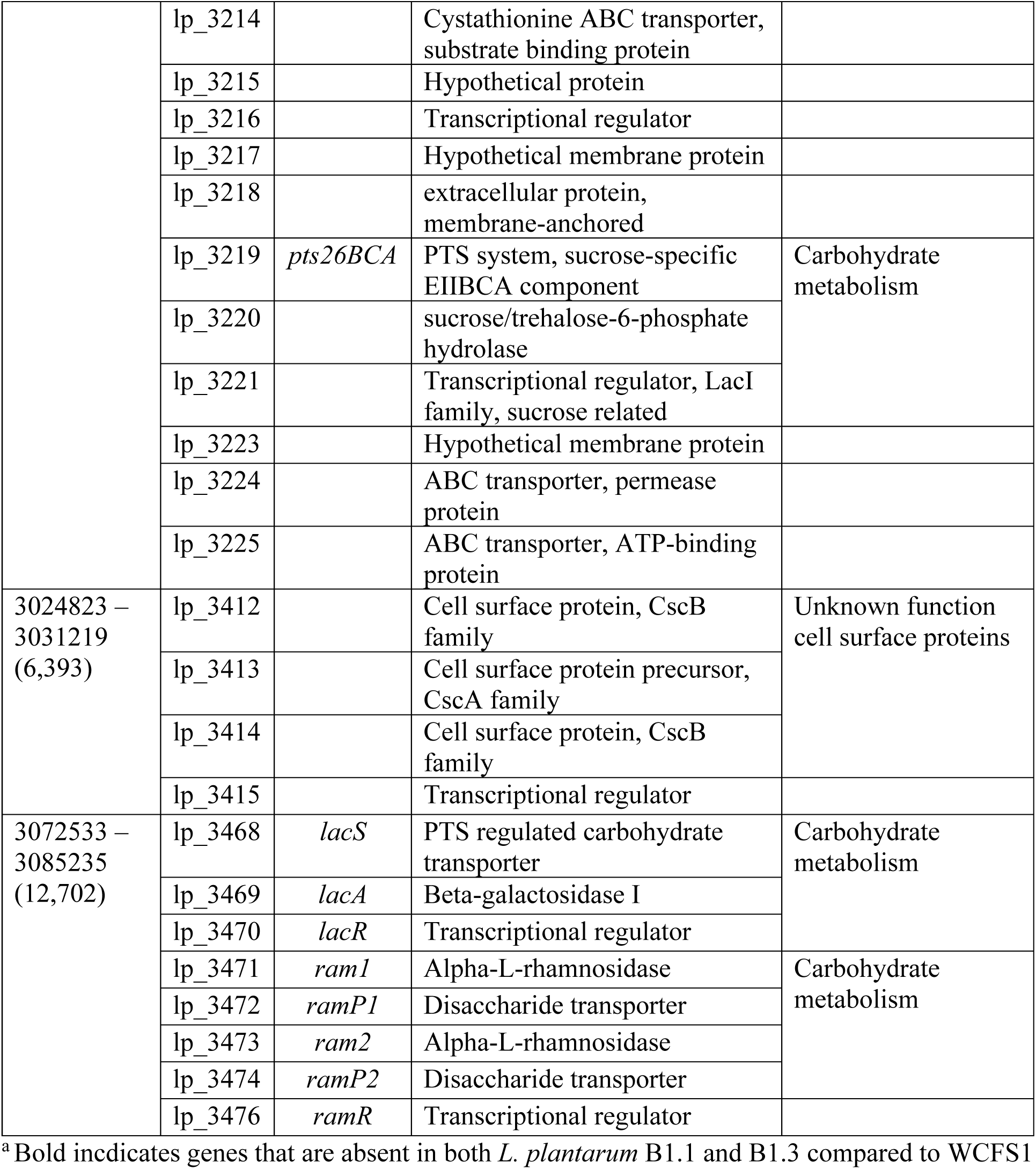
Genes present in *L. plantarum* WCFS1 and absent in *L. plantarum* B1.3.

**Fig. S1.**
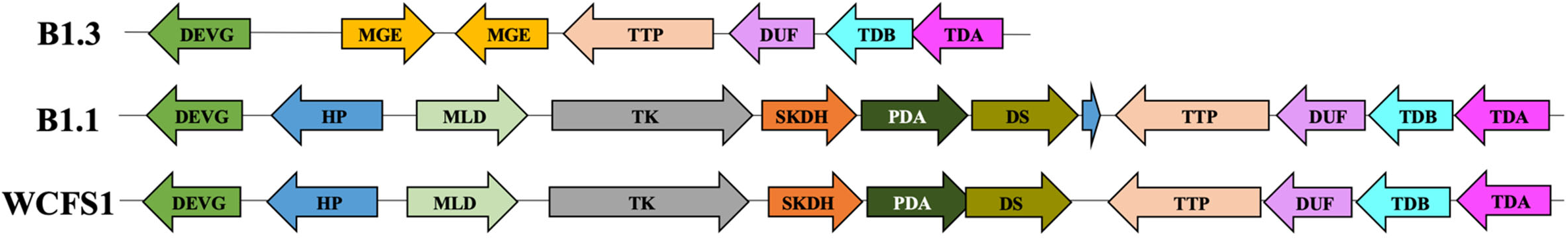
*L. plantarum* B1.3 lacks shikimate pathway genes required for aromatic amino acid biosynthesis. Genome organization of shikimate pathway genes in *L. plantarum* strains. DEVG: fatty acid-binding protein, DegV family (lp_1079), HP: hypothetical protein (lp_1081), MLD: malate/lactate dehydrogenase (lp_1082), TK: transketolase (*tkt2*; lp_1083), SKDH: shikimate 5-dehydrgonase (aroD1; lp_1084), PDA: phosphor-2-deoxyheptonate aldolase/ chroismate mutase (aroA; lp_1085), DS: 3-dehydroquinate synthase (aroB; lp_1086). TTP: tartate transport protein (ttpD; lp_1087), DUF: hypothetical protein, DUF59 family (lp_1088), TDB: L(+)-tartrate dehydrates, subunit B (ttdB; lp_1089), TDA: L(+)-tartrate dehydrates, subunit A (ttdA; lp_1090), MGE: mobile genetic element.

**Fig. S2.**
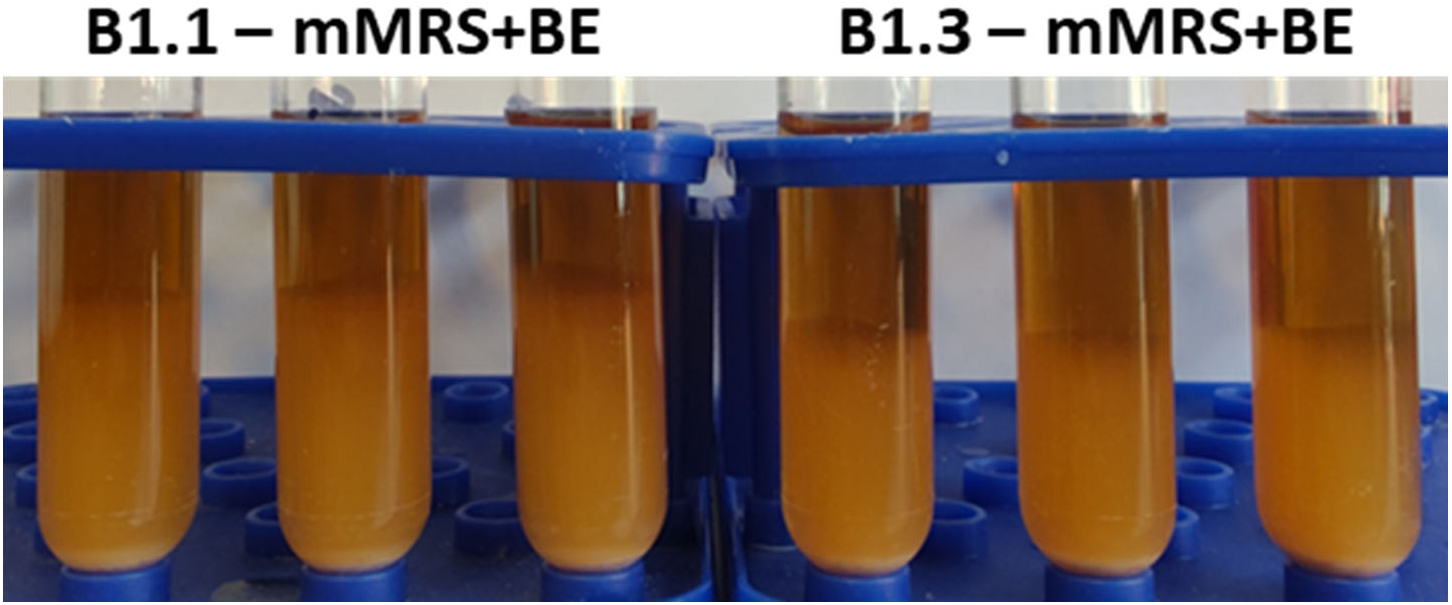
Auto-aggregation of *L. plantarum* B1.1 and B1.3 in mMRS supplemented with beef extract (BE). *L. plantarum* B1.3 and B1.1 cultures were imaged after incubation at 30 °C for 72 h.

**Fig. S3.**
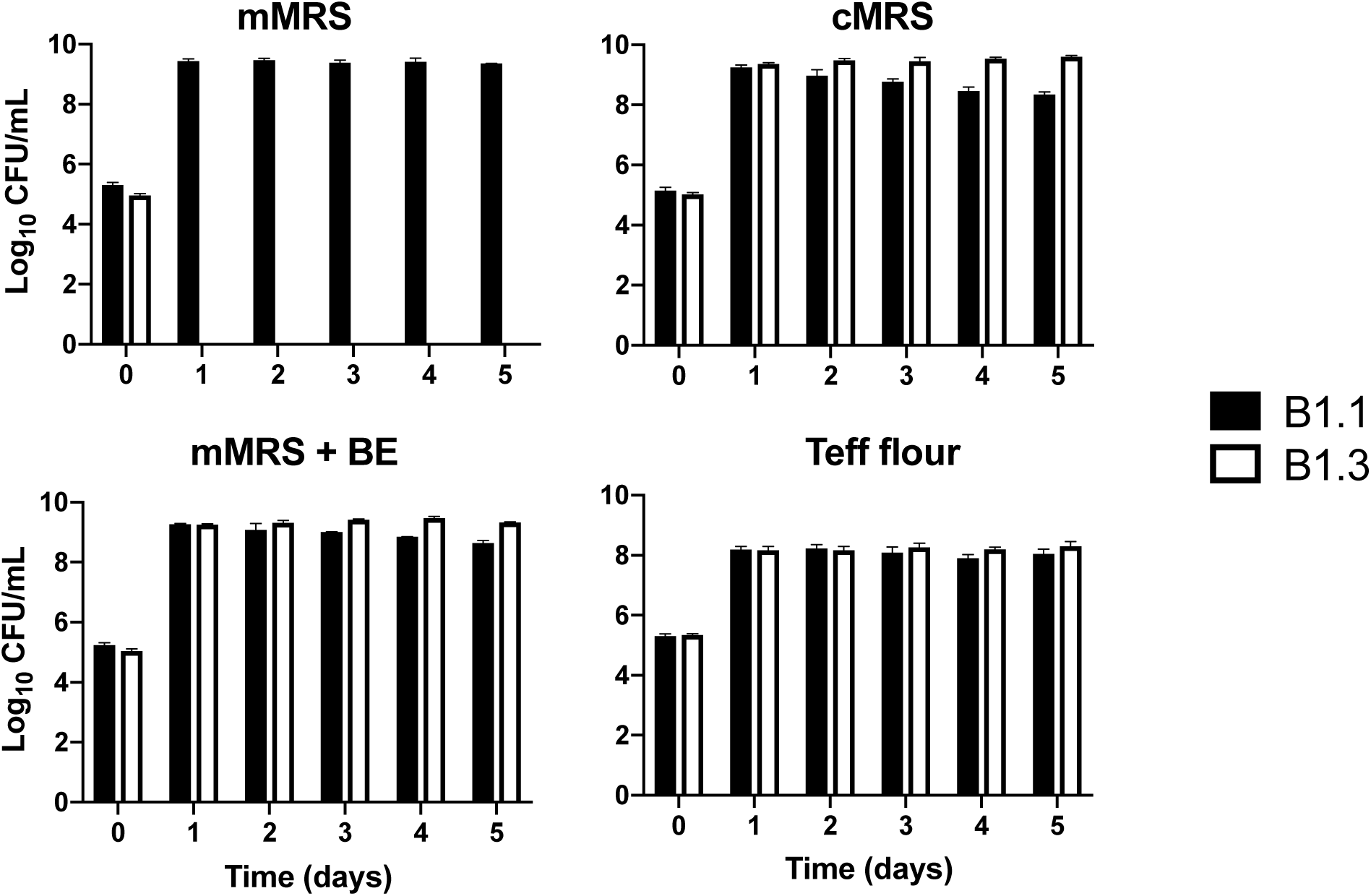
Numbers of *L. plantarum* B1.1 and B1.3 in mMRS, cMRS, mMRS supplemented with beef extract, and a teff flour suspension. Equal numbers of *L. plantarum* B1.1 and B1.3 (10^5^ CFU/ml) were co-inoculated in mMRS, cMRS, mMRS supplemented with beef extract (8 g/L), or teff flour mixed with PBS and incubated at 30 °C for 24 h. A total of 50 µl was transferred into fresh medium (constituting 1% of the final volume) on each of the subsequent five days. The avg ± stdev of three replicate cultures are shown.

**Fig. S4.**
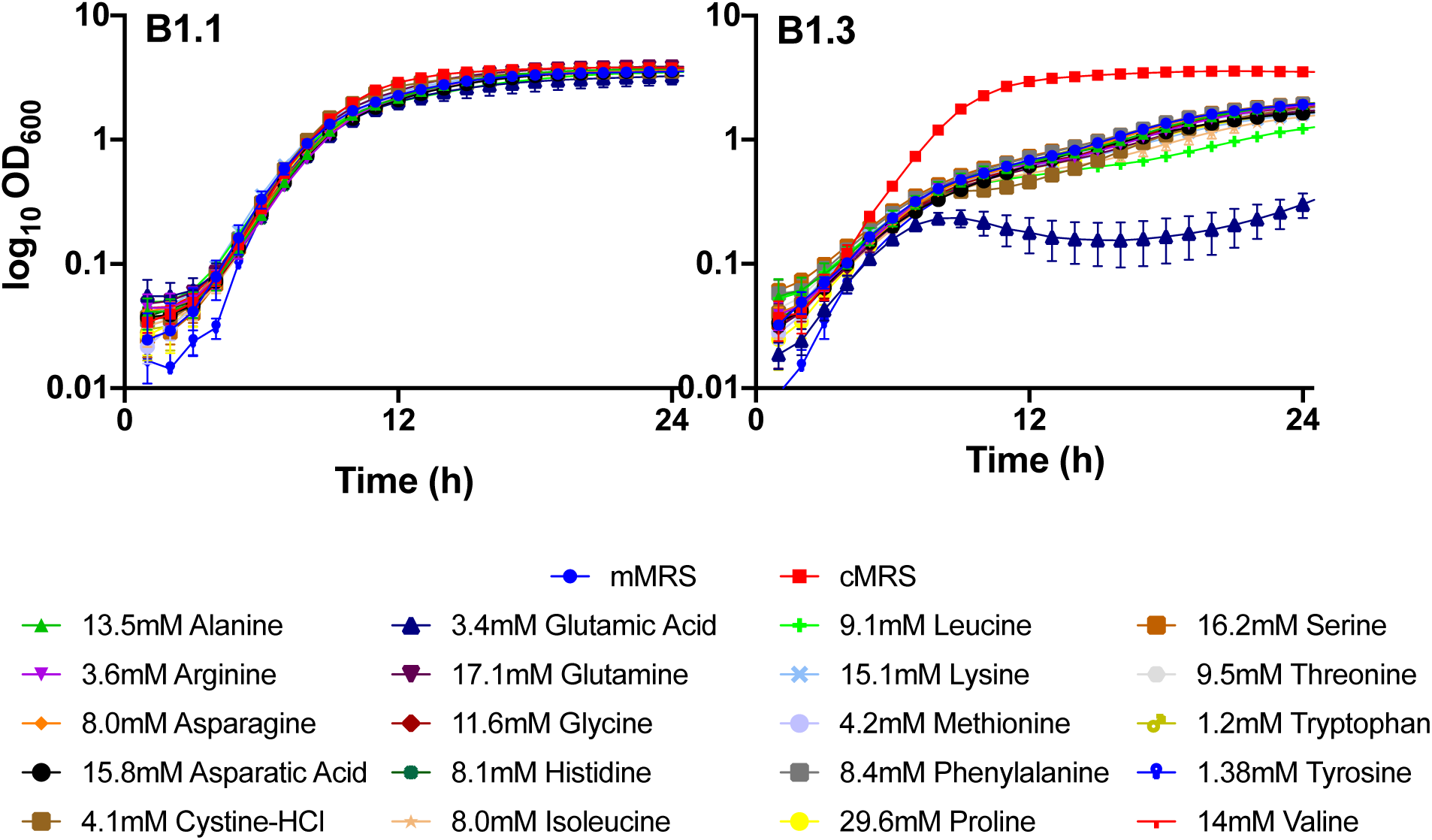
Growth of *L. plantarum* B1.1 and B1.3 in mMRS supplemented with different amino acids. *L. plantarum* B1.1 and B1.3 were inoculated into mMRS supplemented with single amino acids and incubated at 30 °C for 24 h. The avg ± stdev OD_600_ of three replicate wells are shown.

**Fig. S5.**
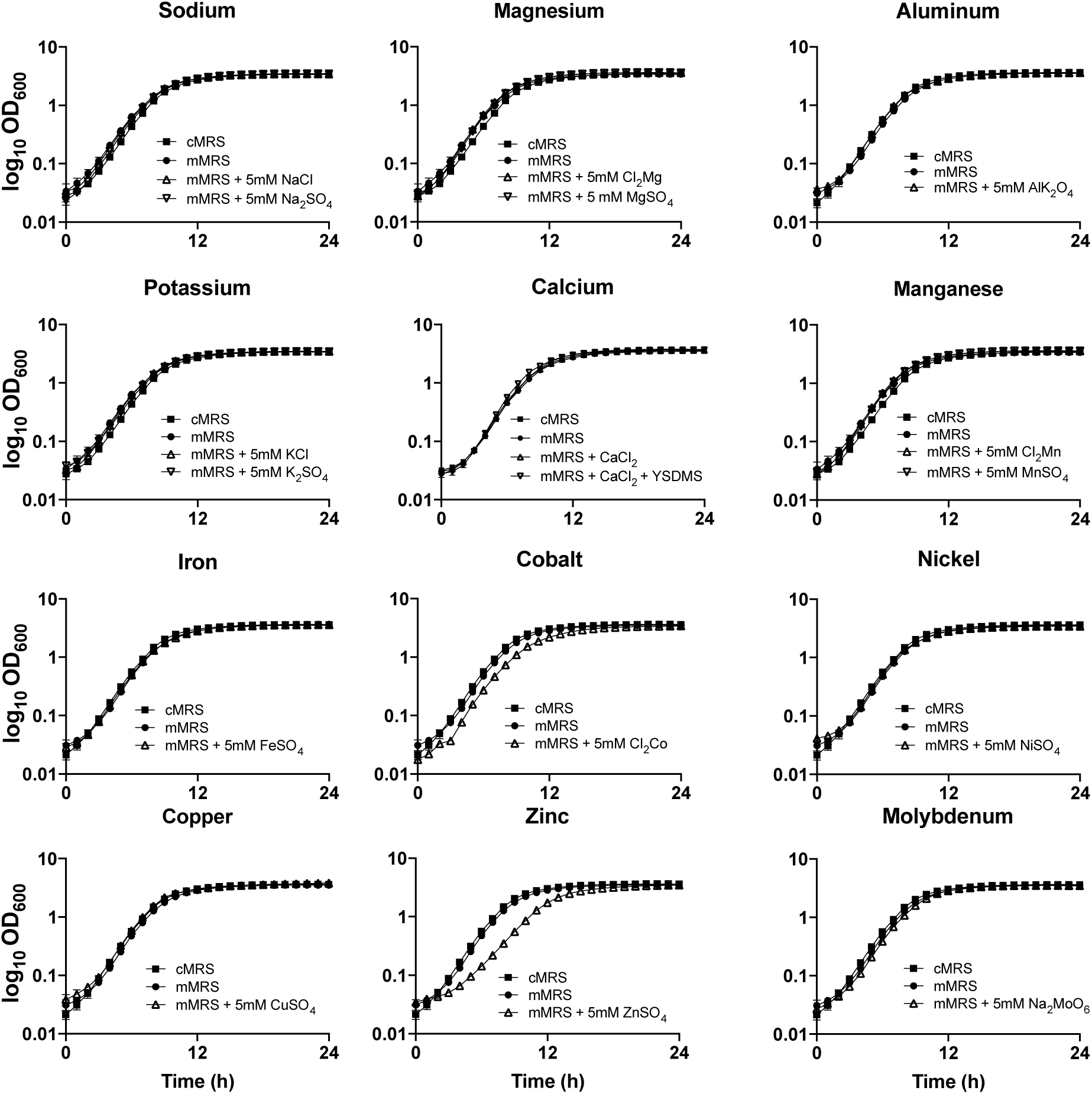
Growth of *L. plantarum* B1.1 in mMRS supplemented trace metals. *L. plantarum* B1.1 were inoculated into mMRS containing 5mM of individual trace metals and incubated at 30 °C for 24 h. The avg ± stdev OD_600_ of three replicate wells are shown.

**Fig. S6.**
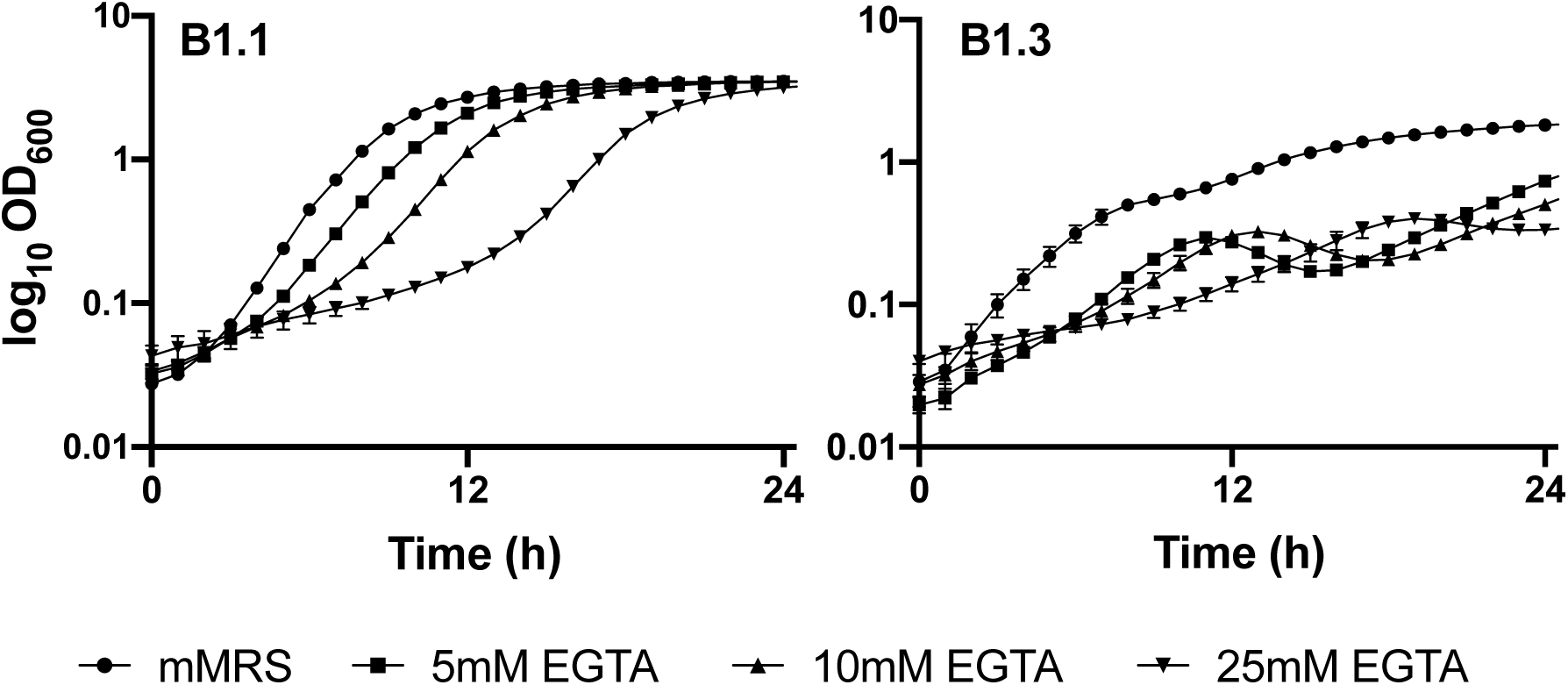
Growth of *L. plantarum* in mMRS supplemented with different concentrations of EGTA. *L. plantarum* B1.1 and B1.3 were incubated at 30 °C for 24 h. The avg ± stdev OD_600_ of three replicate wells are shown.

**Fig. S7.**
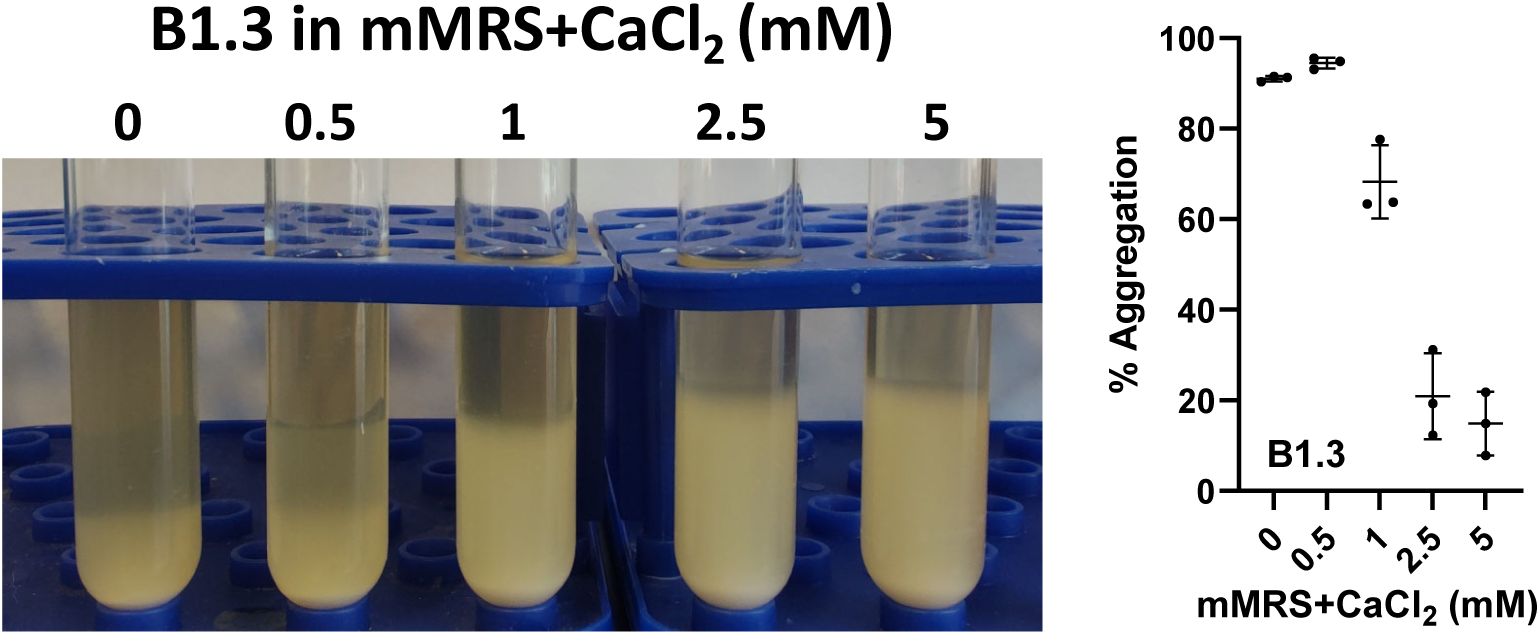
Auto-aggregation of *L. plantarum* B1.3 in mMRS supplemented with different concentrations of CaCl_2_. *L. plantarum* B1.3 cultures were imaged after incubation at 30 °C for 72h. % Aggregation shown as Mean ± SD between triplicates.

**Fig. S8.**
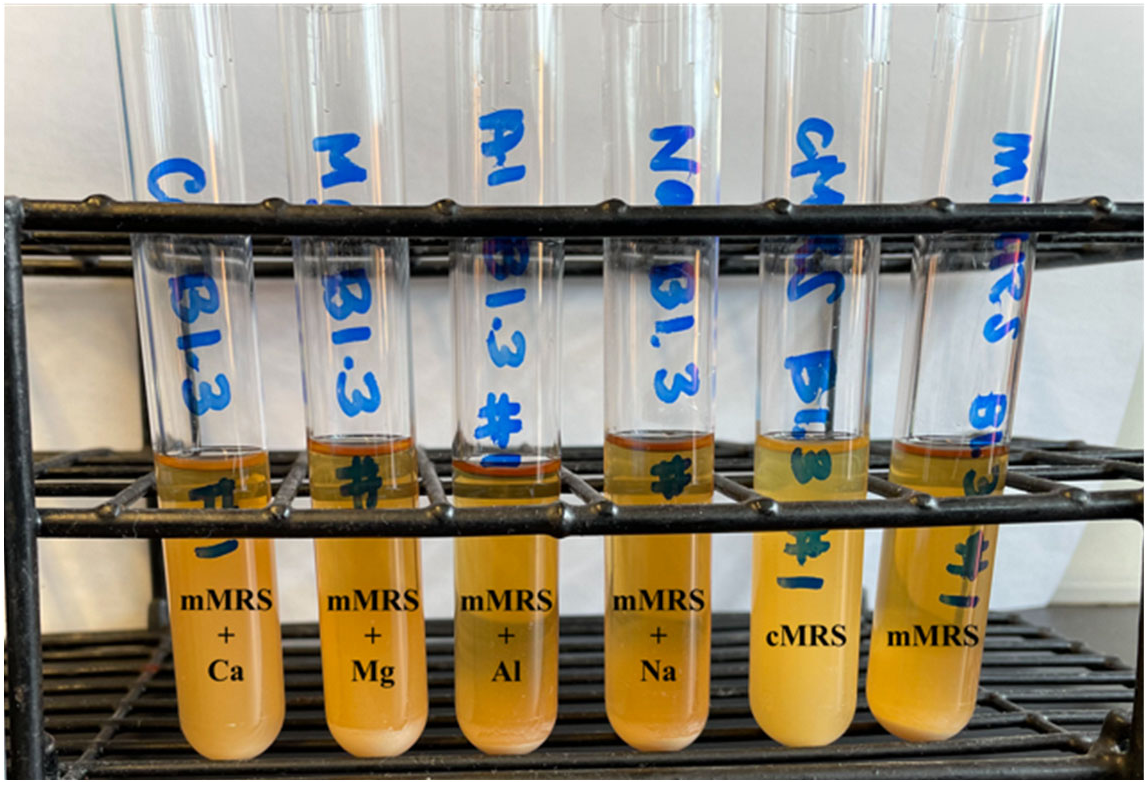
Auto-aggregation of *L. plantarum* B1.3 in mMRS, cMRS, and mMRS supplemented with 5mM of CaCl_2_, MgSO_4_, AlK_2_O_4_, or NaCl. *L. plantarum* B1.3 cultures were imaged after incubation at 30 °C for 72 h.

**Fig. S9.**
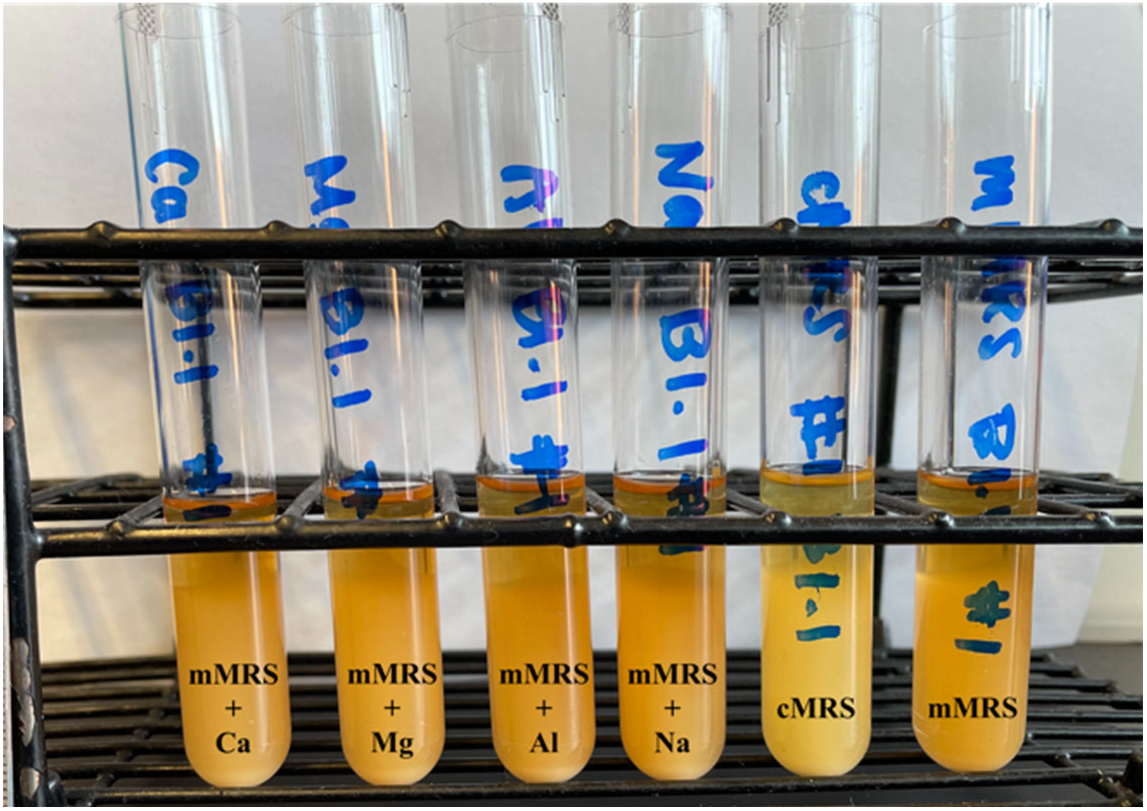
Auto-aggregation of *L. plantarum* B1.1 in mMRS, cMRS, and mMRS supplemented with 5mM of CaCl_2_, MgSO_4_, AlK_2_O_4_, or NaCl. *L. plantarum* B1.1 cultures were imaged after incubation at 30 °C for 72 h.

**Fig. S10.**
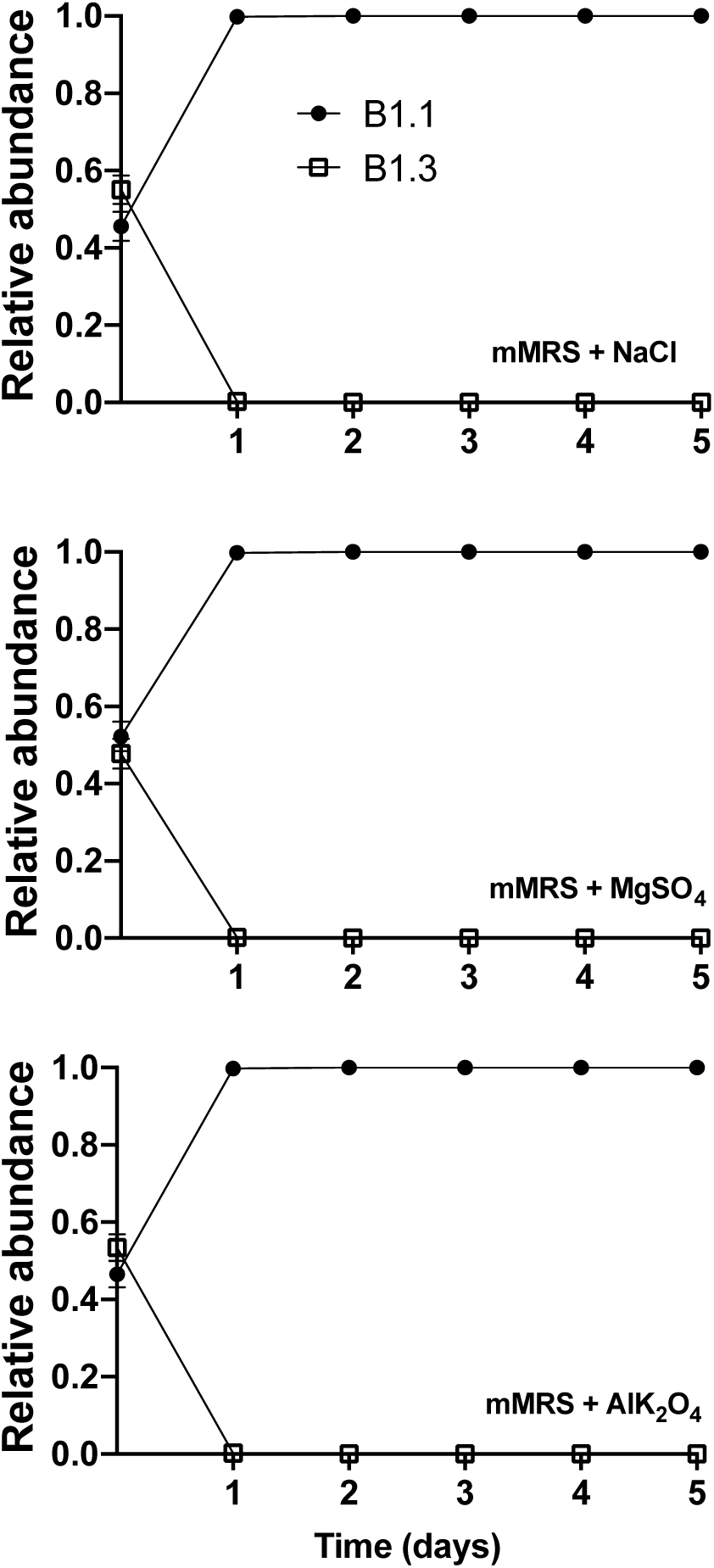
Competitive fitness of *L. plantarum* B1.1 and B1.3 in mMRS 5mM of NaCl, MgSO_4,_ or AlK_2_O_4_. Equal numbers of *L. plantarum* B1.1 and B1.3 (10^5^ CFU/ml) were co-inoculated in mMRS 5mM of NaCl, MgSO_4,_ or AlK_2_O_4_. A total of 50 µl was transferred into fresh medium (constituting 1% of the final volume) on each of the subsequent five days. The avg ± stdev of three replicate cultures is shown.

